# Rejuvenation of the Aged Cerebrovascular System via Protein Corona–Guided Fusogenic Liposome Delivery

**DOI:** 10.64898/2026.03.05.709925

**Authors:** Santny Shanmugarama, Till Gronemann, Boglarka Csik, Roland Patai, Ádám Nyúl-Tóth, Dorina Nagy, Laszlo Hricisak, Mark Nagykaldi, Madison Sanford, Raghavendra Y Nagaraja, Rafal Gulej, Rebeka Kristof, Kiana Vali Kordestan, Evelyn G Brunner, Sharon Negri, Hassan Abushukair, Woncheol Jung, Stefano Tarantini, Siva Sai Chandragiri, Parikshat Sirpal, Shannon Conley, Peter Mukli, Andriy Yabluchanskiy, Priyabrata Mukherjee, Sabrina Berkamp, Nils Hersch, Maithreyan Kuppusamy, Carsten Sachse, Pitter Huesgen, Rudolf Merkel, Tamas Kiss, Zoltan Benyo, Tae Gyu Oh, Zoltan Ungvari, Agnes Csiszar, Anna Csiszar

**Affiliations:** Vascular Cognitive Impairment, Neurodegeneration, and Healthy Brain Aging Program, Department of Neurosurgery, University of Oklahoma Health Sciences Center, Oklahoma City, OK, USA; Oklahoma Center for Geroscience and Healthy Brain Aging, University of Oklahoma Health Sciences Center, Oklahoma City, OK, USA; Forschungszentrum Jülich GmbH, Institute of Biological Information Processing: Mechanobiology (IBI-2), Jülich, Germany; International Training Program in Geroscience, Doctoral School of Basic and Translational Medicine/Department of Public Health, Semmelweis University, Budapest, Hungary; Cerebrovascular and Neurocognitive Disorders Research Group, Hungarian Research Network, Semmelweis University (HUN-REN-SU), Budapest, Hungary; Department of Oncology Science, University of Oklahoma Health Sciences Center, Oklahoma City, OK, USA; The Peggy and Charles Stephenson Cancer Center, University of Oklahoma Health Sciences Center, Oklahoma City, OK, USA; Forschungszentrum Jülich GmbH, Ernst Ruska-Centre for Microscopy and Spectroscopy with Electrons: Structural Biology (ERC-3), Jülich, Germany; Forschungszentrum Jülich GmbH, Central Institute of Engineering, Electronics and Analytics: Analytics (ZEA-3), Jülich, Germany; Faculty of Biology and CIBSS-Centre for Biological Signaling Studies, University of Freiburg, Freiburg, Germany

**Keywords:** ApoE-targeting, fusogenic liposome, resveratrol delivery, cerebrovascular aging, improving blood-brain barrier functions, neurovascular coupling

## Abstract

Brain vascular aging is increasingly recognized as a critical therapeutic target for age-related cognitive decline. Oxidative stress, bioenergetic dysfunction, and molecular damage play central roles in the progression of vascular aging, contributing to cerebrovascular dysfunction and impaired cognitive function. While naturally occurring polyphenols such as resveratrol (RSV) have demonstrated potential in mitigating aging-related pathologies, their poor bioavailability and limited brain targeting efficiency significantly constrain their therapeutic impact. As a result, high doses or advanced drug delivery strategies are necessary to achieve meaningful physiological effects.

We introduce a novel nanocarrier system designed to enhance RSV delivery to the cerebral endothelium by leveraging the natural formation of an apolipoprotein E (ApoE)-enriched protein corona around fusogenic liposomes (FL) *in vivo*. These nanoparticles directly fuse with cytoplasmic cell membranes and thus evade endocytosis. We found that once in the circulation FL spontaneously acquire a protein corona, which is highly enriched in ApoE, a key ligand for brain endothelial low-density lipoprotein receptors (LDLR). Based on this observation, we engineered an ApoE-functionalized protein corona around FL (ApoE-FL) to systematically evaluate whether this mechanism could be exploited for targeted brain delivery. Following optimization and physicochemical characterization, the RSV-loaded liposomes were evaluated *in vitro* using human cerebral microvascular endothelial cells and *in vivo* C57BL/6 aged mice to assess their therapeutic potential. Both FL and engineered ApoE-FL liposomal delivery systems exhibited a strong affinity for endothelial cell membranes *in vitro*. The knockdown of the ApoE receptor, low-density lipoprotein receptor-related protein 1 (LRP1), significantly reduced liposomal docking. Microscopy analysis revealed that both ApoE-FL and non-functionalized FL directly fused with endothelial plasma membranes, thus bypassing intracellular organelles and minimizing lysosomal degradation. This suggests that the naturally formed ApoE corona *in vivo* may contribute to efficient cerebrovascular targeting, a property successfully replicated by the engineered ApoE corona strategy.

*In vivo* biodistribution and kinetic studies demonstrated that especially ApoE-FL achieved enhanced brain-targeting efficiency, prolonged cerebrovascular retention, and extended targeting distance along the arteriovenous axis. This emphasizes that fusogenic liposomes effectively engage almost the entire microvascular network, including capillaries and post-capillary venules. Functionally, fusogenic liposome-delivered RSV improved blood-brain barrier (BBB) integrity, enhanced neurovascular coupling (NVC) responses, and promoted brain vascularization in aged mice. Single-cell RNA sequencing (scRNA-seq) revealed enhanced endothelial angiogenesis and barrier protective transcriptional profiles in cerebrovascular cells treated with ApoE-FL/RSV, suggesting a molecular basis for the observed vascular benefits. Liposomal RSV delivery achieved near-complete cerebrovascular and cognitive rejuvenation in aged mice applying a 2000-fold lower RSV dose than oral administration used as control sample. Thus, ApoE-FL liposomes exhibited exceptionally high delivery efficiency in deeper brain regions, further expanding their therapeutic potential. These findings underscore the importance of targeted drug delivery in optimizing therapeutic outcomes and establish ApoE-functionalized fusogenic liposomes as a promising strategy for mitigating brain vascular aging and cognitive decline.

**Graphical Abstract:** 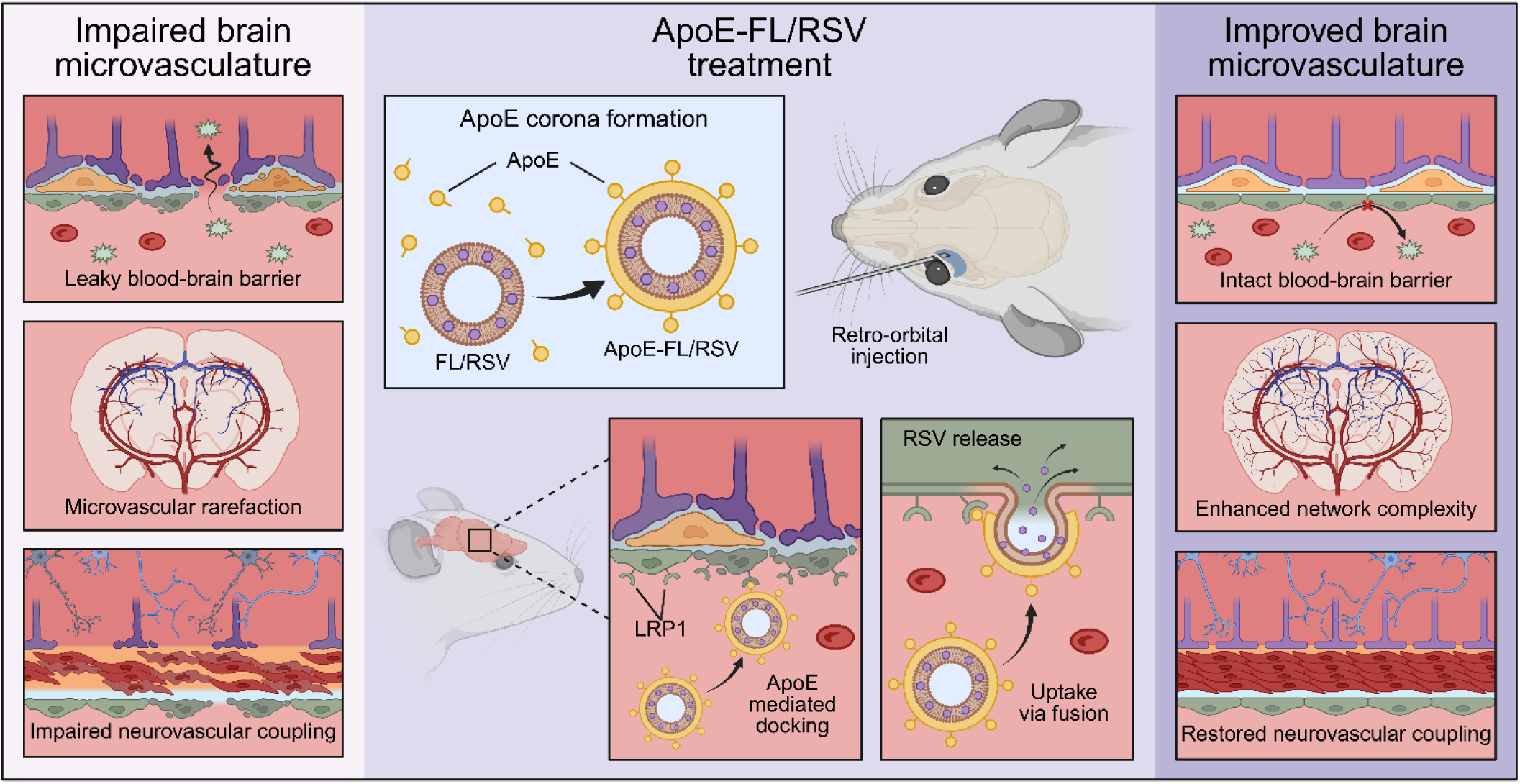

## Introduction

Aging populations worldwide face an increasing burden of vascular cognitive impairment and dementia (VCID), conditions that significantly affect quality of life and place a substantial strain on healthcare systems. VCID, driven by cerebrovascular dysfunction, underscores the intricate relationship between vascular health and cognitive function^1–8^. By targeting the underlying mechanisms of vascular aging, such as endothelial dysfunction and blood-brain barrier disruption, novel therapeutic approaches could not only improve vascular health but also preserve cognitive function. This dual focus offers a promising pathway to mitigate the progression of VCID and enhance overall brain health in aging individuals, providing hope for effective interventions^2, 9–18^. To translate these insights into effective therapies, innovative drug delivery systems are essential to specifically treat the cerebrovascular system. Targeted delivery of therapeutics directly to brain endothelial cells can pave the way for interventions that address both vascular and cognitive health. In this context, the use of resveratrol (RSV) encapsulated in fusogenic liposomes represents a groundbreaking strategy, leveraging the compound’s well-established vascular benefits^19–28^ while addressing the specific needs of the vasculature in the aging brain.

Resveratrol, a naturally occurring polyphenolic compound, has well-documented benefits for aging vasculature^23, 27, 29^. It improves endothelial function by reducing oxidative stress, suppressing inflammation, and activating key mitochondrial molecular pathways such as Sirtuin 1 (SIRT1), AMP-activated protein kinase (AMPK), and endothelial nitric oxide synthase (eNOS)^19, 21, 22, 24, 25, 28, 30, 31^. These mechanisms collectively enhance vasodilation^27^, decrease vascular stiffness, and protect against atherosclerosis in the large vessels, indirectly supporting neuronal health by improving cerebral blood flow (CBF)^20, 26^. However, the therapeutic potential of RSV in the microcirculation, which undergoes some of the most significant age-related changes^32^, is limited by its poor bioavailability, rapid metabolism, and low systemic stability^33^. These factors result in inadequate drug concentrations reaching target tissues such as the brain, ultimately reducing its effectiveness^34^.

In order to increase RSV’s bioavailability several cationic liposomal formulations have been developed in the past^35–37^. Their cellular uptake route was mainly found to be clathrin- or caveolae-mediated endocytosis^37^. However, the efficiency of endocytic uptake is limited^38^. Our previous studies demonstrated that the presence of an aromatic molecule in cationic lipid nanoparticles completely changes the uptake route from endocytosis to fusion with the cytoplasmic membrane^13,14^. This way endocytic pathways are evaded resulting in significantly reduced degradation of the encapsulated compound and strongly increased cytoplasmic concentrations. For polyphenols such as RSV, which are characterized by poor bioavailability and rapid systemic clearance, fusogenic liposomes offer critical advantages. Their bilayer structure protects RSV from enzymatic degradation and metabolic inactivation, ensuring sufficient stability and prolonged circulation. We have successfully tested RSV payload delivery by FL *in vitro*^21^ as well as *in vivo*^40^. These findings confirm that RSV reaches the cerebral vasculature via FL-mediated transport. However, while these effects are promising, they remain modest and variable, suggesting that RSV delivery is not yet optimally targeted or efficient within the brain. To fully harness the therapeutic potential of RSV, particularly in the context of age-related cerebrovascular dysfunction, an enhanced delivery strategy with increased brain specificity is needed.

To keep the targeting concept as straightforward as possible, we exploited a naturally occurring phenomenon. Upon interaction with biological fluids like serum, nanoparticles including fusogenic liposomes are enveloped by a protein corona, a layer of proteins and other biomolecules that adsorb onto their surface^41^. We found that the spontaneously formed protein corona around fusogenic liposomes is primarily composed of apolipoproteins, such as ApoE, ApoA, and ApoB, along with other serum factors. The presence of apolipoprotein ApoE is of significant interest due to its well-documented role in interaction with brain endothelial cells and facilitating transport across the BBB^42, 43^. Most liposomal functionalization strategies targeting the brain rely on ApoE-derived peptides rather than the full-length protein, primarily focusing on the ApoE-LRP1 receptor-ligand interaction^42, 44^. However, this approach sacrifices several beneficial cellular functions of the full-length protein, including its role in maintaining BBB integrity^45^.

Functionalizing liposomes with ApoE peptides typically require time- and cost-intensive chemical reactions. In contrast, our findings reveal that full-length ApoE protein spontaneously integrates into the protein corona of fusogenic liposomes. This spontaneous binding eliminates the need for chemical modifications, providing an efficient and accessible method for functionalizing FL loaded with RSV. The resulting particles, enriched with full-length ApoE, retain their complete functionality and are effectively targeted to cerebral endothelial cells via LRP1 docking. Such ApoE-functionalized fusogenic liposomes (ApoE-FL) could represent a transformative advance in the treatment of age-related cerebrovascular and neurodegenerative diseases, leveraging the natural pathways of lipid and protein transport for highly efficient drug delivery.

Most experimental nanocarrier systems have been evaluated primarily *in vitro* using cell culture systems, with limited investigation of *in vivo* functional outcomes, where they often fail to achieve comparable drug delivery due to metabolic degradation or impaired targeting^46^. In this study, we comprehensively evaluated the potential of ApoE-functionalized fusogenic liposomes (ApoE-FL/RSV) to achieve precise and efficient delivery of RSV to the aging cerebrovascular system and to promote vascular and cognitive rejuvenation. We aimed to optimize brain-specific targeting and uptake of RSV-loaded liposomes following intravenous administration in both young (3–9-month-old) and aged (18–25-month-old) C57BL/6 mice. We evaluated whether full-length ApoE functionalization enhances selective brain delivery and improves cerebrovascular structure and function in aged animals. Finally, we assessed whether these vascular effects translate into measurable cognitive benefits, thereby defining the contribution of ApoE-FL/RSV delivery to maintaining neurovascular health during aging.

## Results

### Formulation and characterization of ApoE-FL/RSV and FL/RSV liposomes

In our previous study, we established the preparation of fusogenic liposomes loaded with RSV (FL/RSV) and demonstrated their beneficial effects on neurovascular coupling in aged C57BL6 mice *in vivo*, while confirming that empty FLs have no observable effects (for particle characterization, see Extended Data Fig. 1)^40^. When liposomes are injected into the bloodstream, serum proteins interact with the drug delivery vehicles, forming a passive protein corona on the liposomés surface^41^. As the protein corona fully envelops the particles, its formation often masks the original material properties, giving rise to a new, biologically active identity. (Fig. 1A). We analyzed the composition of the protein corona on the FL/RSV particles after 1-hour *in vitro* incubation with human or mouse plasma (Fig. 1B) using liquid chromatography-tandem mass spectrometry (LC-MS/MS). Our results revealed reduced proportion of immunoglobulins and serum albumin in the protein corona compared to the composition of reference plasma samples, while the proportion of apolipoproteins was significantly increased (Fig. 1B). Apolipoproteins, in general, are characterized by a strong intrinsic tendency for the formation of nanoscopic lipid structures based on their high binding affinity to phospholipids, triacylglycerides, and cholesterol in the blood plasma^47^. Notably, ApoE, a member of the apolipoprotein family, plays a crucial role in targeting lipid complexes in plasma or interstitial fluids to specific cell-surface receptors. Among these, LRP1 stands out for its high expression on cerebrovascular endothelial cells, enabling precise interaction and uptake within the brain vasculature^48^.

**Figure 1.**
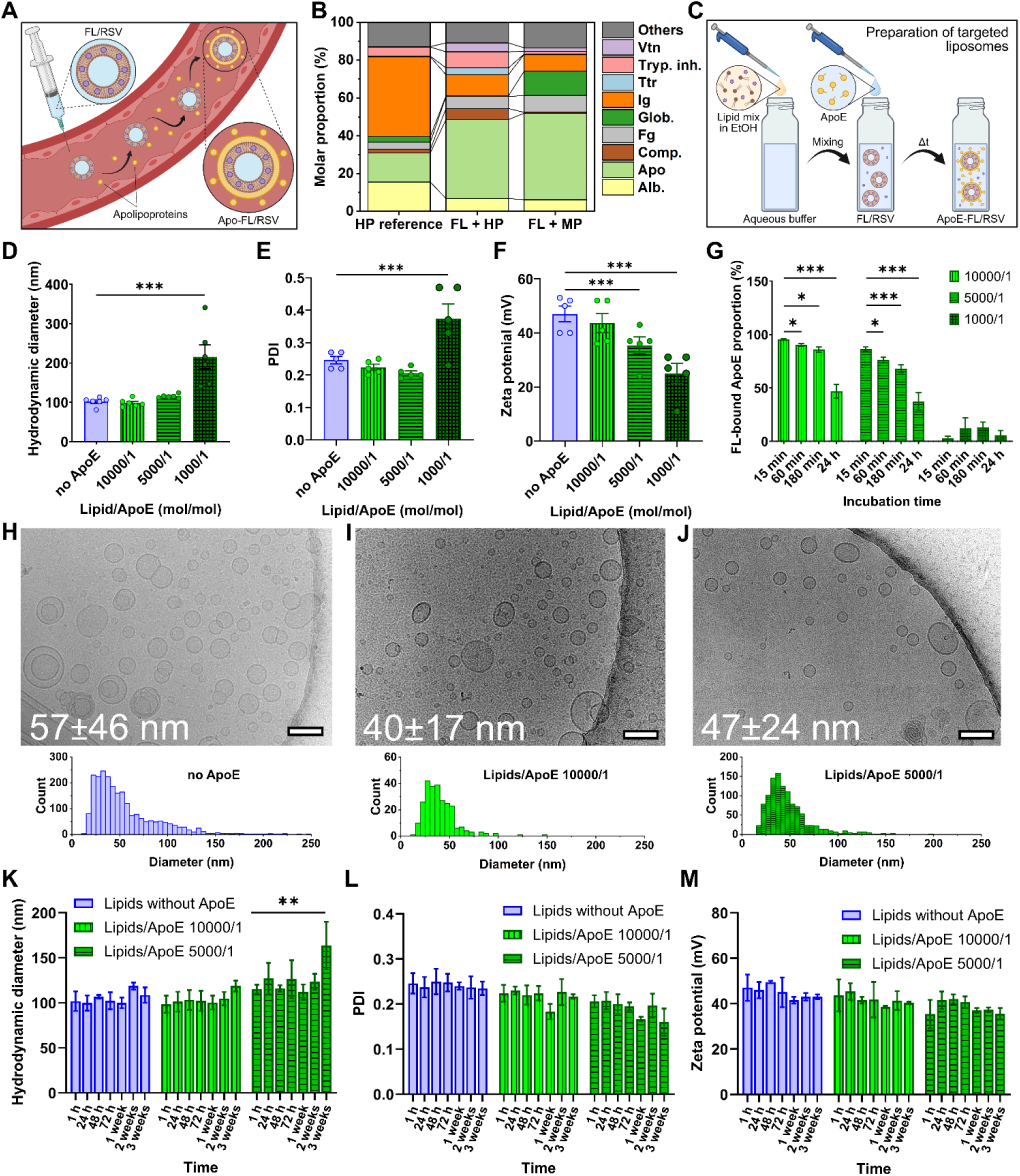
Formulation and physical characteristics of fusogenic liposomes enhanced with an ApoE protein corona. **A)** When FL are intravenously injected, a specific corona composed of different plasma proteins forms on the drug delivery particle surface. **B)** LC-MS/MS-based compositional corona analysis of FL incubated in human (HP) or mouse plasma (MP). Protein corona composition on the FL surface is compared with that of fresh HP samples. **C)** RSV-loaded FL were prepared using the ethanol-injection method. The formed particles were subsequently incubated in a human ApoE solution. Based on the high binding affinity of ApoE to lipid particles, ApoE-coating of FL/RSV liposomes has been assumed. **D)** Hydrodynamic diameter, **E)** particle homogeneity, characterized by the polydispersity index (PDI), and **F)** zeta potential of ApoE-coated FL/RSV (2/0.4 mol/mol) were determined via dynamic and electrophoretic light scattering, respectively. Lipid/ApoE ratio was varied between 10000/1 and 1000/1. Liposomes without ApoE served as control. Zeta potential decreased significantly when ApoE attached to the particle surface and screened the positive lipid charge while hydrodynamic diameter as well as PDI value increased only at a lipid/protein molar ratio of 1000/1 mol/mol indicating inhomogeneous particle formation. **G)** Analysis of ApoE-FL complex stability in blood plasma over time indicates high stability in case of low ApoE amounts (10000/1 and 5000/1 lipid/ApoE ratio). **H)** Representative cryo-TEM micrographs of FL/RSV (2/0.4 mol/mol) and **I-J)** ApoE-coated FL/RSV particles clearly show the presence of unilamellar liposomes. Size distribution curves are depicted below. Average particle diameters and standard deviations of the distributions are indicated. Scale bars correspond to 100 nm. Particles appear stable over a period of 3 weeks based on the monitoring of characteristic physicochemical parameters such as **K)** hydrodynamic diameter, **L)** PDI, and **M)** zeta potential. All data are shown as mean±SEM (n≥5). Statistical significance was calculated using the two-sample t-test and is indicated by *p<0.05, **p<0.01, ***p<0.001.

Given the strong binding affinity of full-length ApoE to lipid particles, we tested whether the spontaneous binding of ApoE to FL eliminates the need for chemical modifications, providing an efficient and accessible method for functionalizing FL loaded with RSV (Fig. 1C). Therefore, human ApoE has been chosen and coupled to the FL/RSV surface via incubation of the full-length protein with the drug-loaded particles. Selecting the optimal ApoE-to-lipid molar ratio is crucial for maintaining the stability and functionality of ApoE-functionalized fusogenic liposomes (ApoE-FL/RSV). At ApoE/lipid molar ratios between 1/10000 and 1/5000, the hydrodynamic diameter (98 ± 9 nm and 115 ± 5 nm (Fig. 1D) and polydispersity index (PDI 0.22 ± 0.02 and 0.21 ± 0.02, respectively) (Fig. 1E) remained stable, indicating uniform and well-formed particles. However, when the ratio exceeded 1/1,000, the hydrodynamic diameter (215 nm ± 74 nm), and PDI (0.37± 0.1) increased significantly, suggesting particle aggregation at higher ApoE concentrations. Concurrently, zeta potential decreased progressively, from 47 mV ± 6 mV to 25 ± 8 mV, reflecting increased ApoE coverage on the liposome surface (Fig. 1F). In addition to signs of particle aggregation, samples with high ApoE-to-lipid ratios (1/1000) showed a marked decrease in complex stability. Stability studies of ApoE-FL in the presence of full plasma revealed that only about 10% of ApoE remained associated with the liposomal surface (Fig. 1G), with the majority either washed off or displaced by other plasma proteins. In contrast, at lower ApoE-to-lipid ratios of 1/10,000 and 1/5,000, a significantly higher proportion of ApoE remained bound to the liposomes, exceeding 90% and 80%, respectively, within the first hour.

Cryogenic transmission electron microscopy (Cryo-TEM) analysis further confirmed that at these optimal ratios, the particles were uniformly distributed, forming unilamellar liposomes with an average core diameter of approximately 50 nm (Fig. 1H–J), indicating excellent structural integrity and uniformity. The observed difference between the hydrodynamic diameter measured by dynamic light scattering (DLS) and the physical diameter determined by cryo-TEM reflects the distinct properties captured by each technique. The hydrodynamic diameter represents how particles behave in solutions, incorporating contributions from the hydration shell, surface-bound molecules, and particle dynamics. In contrast, cryo-TEM measures the actual physical size of the particle core. In addition, DLS disproportionately weighs larger particles in the sample. Consequently, hydrodynamic diameters are typically larger than diameters obtained by cryo-TEM.

Furthermore, long-term storage studies showed that ApoE-FL/RSV particles remained stable for up to 2 weeks at 4 °C, with no significant changes in their physicochemical properties (Fig. 1K–M). Based on these findings, we selected an ApoE-to-lipid ratio of 1/5,000 mol/mol, as ratios within the 1/10,000 to 1/5,000 range provided optimal particle size, uniformity, and stability of the complex, whereas higher ratios compromised these properties.

To identify the RSV concentration that can be effectively encapsulated in FL without altering the liposomal physicochemical properties, we tested RSV concentrations ranging from a 2/0.2 to a 2/0.8 mol/mol lipid/RSV ratio. By measuring hydrodynamic diameter and zeta potential of the FL/RSV complexes (Extended Data Fig. 1B–C), we found that the incorporation of RSV in the investigated concentration range did not alter these parameters. While RSV encapsulation into FL was found to be feasible even at high concentrations, it must be considered that the use of excessive drug concentrations could trigger toxic effects. Our previous study demonstrated sufficient efficacy *in vivo* at a lipid/RSV ratio of 2/0.4 mol/mol with no toxicity^49^. Therefore, we used this RSV concentration for all *in vitro* and *in vivo* functional experiments. This corresponds to an RSV dose 2000-fold lower than that usually applied in oral administration^19^ and was used in our *in vivo* control studies.

### FL/RSV and ApoE-FL/RSV cellular uptake via fusion and subcellular distribution in hCMECs

The *in vitro* endothelial uptake of FL particles, with or without RSV, has been previously tested^49,50, 51^ and identified as a highly efficient cytoplasmic membrane fusion process. This mechanism involves the close interaction of the lipid bilayer of FL particles with the cellular plasma membrane, resulting in complete mixing of the two membrane structures^50, 51^ (Fig. 2A). Importantly, it has been demonstrated that empty FL particles have no effects on cellular function^40^. FL particles loaded with RSV (FL/RSV) achieved a fivefold higher intracellular RSV concentration *in vitro* compared to traditional cationic delivery systems, highlighting their superior efficiency for drug delivery^49^. The newly developed ApoE-functionalized particles feature a surface enriched with ApoE protein, designed to enhance attachment to the cellular plasma membrane by targeting LRP1 receptors on brain endothelial cells (Fig. 2A). While this functionalization improves targeting specificity, the dense protein corona may negatively impact membrane fusion efficiency by acting as a barrier to direct lipid bilayer interaction^52^. This could impair intracellular payload delivery and reduce therapeutic effects. To address these concerns, we tested the cellular uptake of ApoE-FL/RSV particles *in vitro* using human brain endothelial cells (hCMEC/D3) to evaluate whether the benefits of ApoE enrichment outweigh potential fusion limitations. The incorporation of the lipophilic fluorescent dyes 3,3’-dioctadecyloxacarbocyanine perchlorate (DiO), 1,1’-dioctadecyl-3,3,3 ’,3 ’-tetramethylindodicarbocyanine 4-chlorobenzenesulfonate (DiD), or 1,1’-dioctadecyl-3,3,3 ’,3 ’-tetramethylindotricarbocyanine iodide (DiR) from the carbocyanine family into the liposomal membrane, enabled the localization of the delivery particles and the monitoring of their subsequent distribution within cell organelles using fluorescence microscopy.

**Figure 2.**
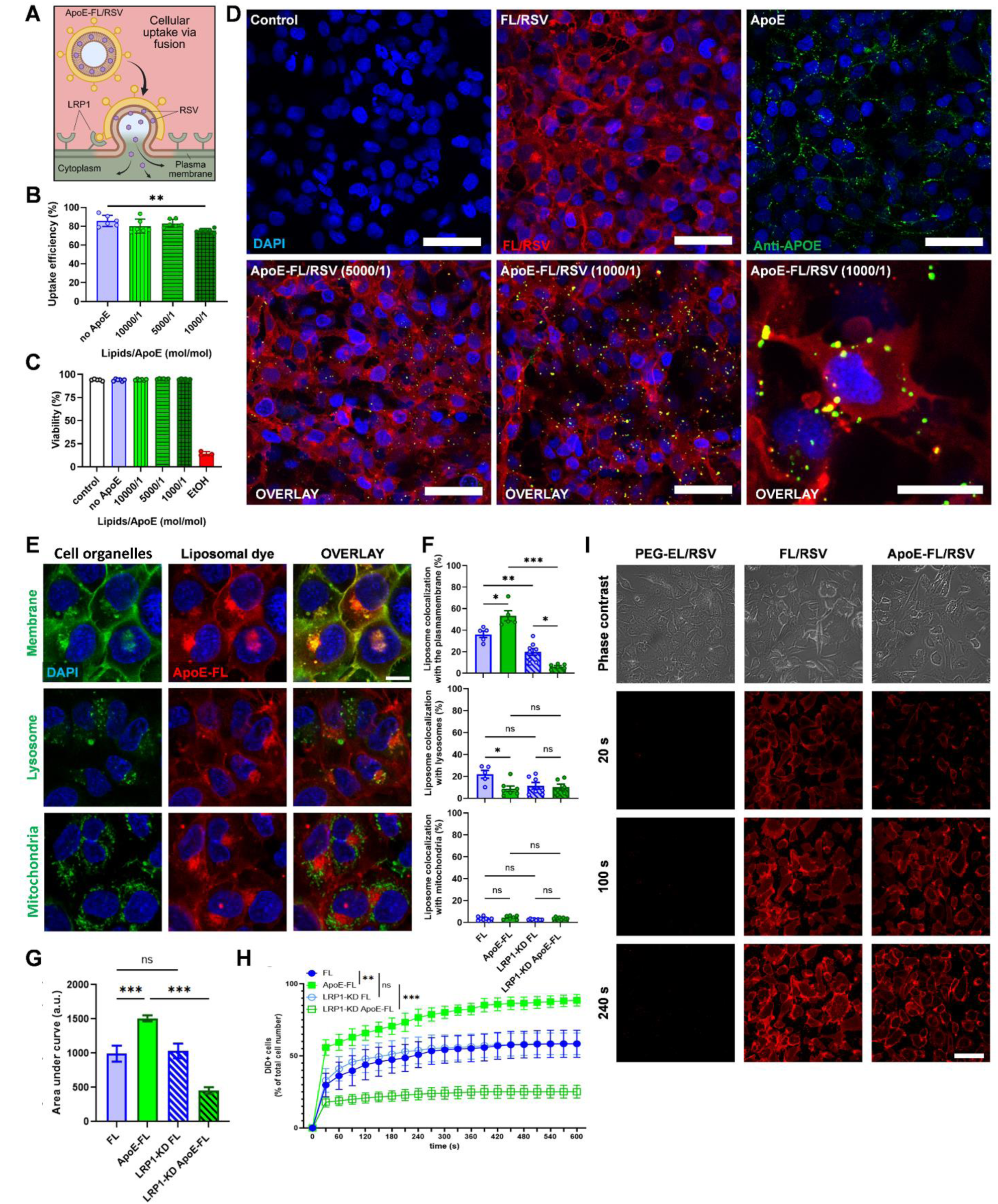
*In vitro* examination of the fusion dynamics of ApoE-FL on human cerebral endothelial cell cultures. **A)** Schematic illustration of the uptake mechanism of FL/RSV with ApoE protein corona. **B)** Uptake efficiency and **C)** viability of hCMECs upon treatment with ApoE-FL/RSV. FL/RSV-treated and non-treated cells were used as control, respectively. **D)** Detection of ApoE on hCMEC/D3 cells fused with ApoE-FL/RSV exhibiting lipid/ApoE ratios of 5000/1 and 1000/1. ApoE presence on the cell surface was proven by a fluorescent anti-ApoE AB (green). Note the increased ApoE signal density in the case of the 1000/1 lipid/ApoE ratio. Untreated cells were used as control. FL/RSV liposomes contained the fluorescent lipid tracer DiR (red) for membrane fusion visualization. Anti-ApoE AB binding was demonstrated by incubating the cells in a highly concentrated ApoE solution. Scale bars correspond to 50 µm and 20 µm for the zoom in. **E)** Distribution of the lipid tracer dye DiD and its colocalization with different cell organelles of hCMEC cells 15 min after ApoE-FL/RSV treatment. Scale bar yields for all, 10 µm. Different cell organelles have been labelled by fluorescent staining in separate cultures. **F)** LRP1 has been knocked down in hCMEC cells to investigate the importance of ApoE in the uptake mechanism. For each organelle, the ratio between DiD/organelle co-staining, and total DiD signal has been quantified. **G)** Uptake efficiency was determined based on the DiD intensity 5 min after treatment with FL/RSV and ApoE-FL/RSV. **H)** To determine uptake kinetics, DiD signal intensity was monitored over a total time period of 10 minutes after treatment. **I)** Representative micrographs of hCMECs at different time points after FL/RSV and ApoE-FL/RSV treatment showing the timeline of the uptake process. A RSV-loaded and PEGylated liposome formulation (PEG-EL/RSV) served as control for no uptake. Scale bars, 50 µm. Again, all data are shown as mean±SEM (n≥5). Statistical significance is indicated by *p<0.05, **p<0.01, ***p<0.001 using one-way analysis of variances (ANOVA) with Tukey’s post-hoc test (F and G), or two-way ANOVA (H).

Following the fluorescence signal of DiR, we found that uptake efficiency was reduced only at a very high ApoE surface concentration (ApoE/lipid ratio of 1/1000 mol/mol) (Fig. 2B), without affecting cell viability (Fig. 2C). Membrane fusion was consistently observed across all conditions, indicated by the uniform fluorescence signal of the DiR lipid tracer in the cellular plasma membrane (Fig. 2D–E). Additionally, ApoE presence on the hCMEC/D3 cell surface was confirmed using a fluorescently labeled anti-ApoE antibody, which appeared as dot-like green signals. A similar pattern was observed when cells were incubated with a highly concentrated ApoE protein solution alone (Fig. 2D). These findings suggest that while a highly concentrated ApoE layer can slightly reduce uptake efficiency, functionalized liposomes in the ApoE/lipid concentration range of 1/10,000-1/5,000 maintain their ability to fuse with the cell membrane and effectively deliver their payload.

Given that functionalization with ApoE and higher ApoE concentration on the liposome surface could negatively impact fusion and shift the uptake mechanism toward LRP1-dependent receptor mediated endocytosis and thereby target subcellular organelles, we specifically aimed to assess these potential changes in fusion dynamics and intracellular distribution (Fig. 2E–F). This analysis was fundamental to determine how ApoE functionalization influences not only cellular uptake but also the intracellular fate of the liposomes, including their potential to avoid lysosomal degradation. As shown in Fig. 2F, ApoE-FL liposomes exhibited markedly higher plasma membrane labeling, with ∼50% of the membrane volume positive for the DiD liposomal tracer dye, indicating enhanced membrane association and fusion efficiency. An LRP1 knockdown cell line was established and the knockdown significantly reduced the membrane co-localization of both FL and ApoE-FL particles; however, the reduction was more pronounced for ApoE-FL, underscoring the importance of ApoE–LRP1 interactions in promoting membrane engagement of ApoE-functionalized liposomes. (For creating LRP1 knockdown (KD) hCMEC cells, see Extended Data Fig. 2 and the Materials and Methods section). Collectively, these findings demonstrate that ApoE functionalization substantially improves plasma membrane targeting through LRP1-dependent mechanisms. Importantly, lysosomal localization was significantly reduced for ApoE-FL liposomes, with only ∼10% of lysosomal volume labeled compared to ∼25% for FL. This finding counters the initial concern that incorporation of ApoE into the protein corona might enhance lysosomal targeting and degradation. LRP1 knockdown did not further alter lysosomal co-localization, indicating that the reduced lysosomal trafficking of ApoE-functionalized liposomes is independent of LRP1. Mitochondrial co-localization with both FL and ApoE-FL signals was minimal, involving less than 4.0–5.5% of mitochondrial volume, suggesting that mitochondria are not a primary target for either FL or ApoE-FL liposomes (Fig. 2E–F).

Our primary concern with ApoE functionalization was that the added protein corona might slow membrane fusion or alter fusion dynamics. However, real-time tracking experiments (Fig. 2I–H) showed that FL and ApoE-FL liposomes displayed comparable fusion kinetics, with both reaching a plateau within approximately 1 minute. LRP1 KD significantly reduced the maximal membrane labeling achieved by ApoE-FL liposomes, although the overall fusion dynamics remained unchanged, as the plateau was still reached within the same time frame. In contrast, the effect of LRP1 KD on FL particles was less pronounced. Blocking the direct contact between plasma membrane and the liposomal surface with the synthetic polymer polyethylene glycol (PEG) prevented liposomal uptake via membrane fusion (Fig. 2I). These findings underscore the critical role of LRP1 specifically in supporting the efficiency, but not the rate of membrane engagement for ApoE-functionalized liposomes (Fig. 2G–H).

To further investigate the cellular distribution of fusogenic liposomes with or without ApoE-targeting, we analyzed dye colocalization with subcellular organelles at 15 minutes, 1 hour, 3 hours, and 24 hours (Extended Data Fig. 3). In both cases, plasma membrane labeling with liposomal DiD plateaued by 3 hours and decreased significantly by 24 hours, whereas endoplasmic reticulum (ER)/Golgi colocalization increased markedly only at the 24-hour time point. This inverse relationship supports the conclusion that liposomes undergo rapid fusion with the plasma membrane, followed by incorporation into the ER/Golgi via slower, natural membrane turnover, rather than through the energy-intensive process of endocytic uptake, which takes approximately 30 min on endothelial cells in the BBB^53^. Lysosomal colocalization increased up to 3 hours but stabilized at a minimal level of less than 10%. These findings provide further evidence of efficient liposome fusion with the cytoplasmic membrane.

### RSV delivered via ApoE-FL and FL effectively reduces oxidative stress in hCMECs and restores cellular viability

FL encapsulated drug follows a different intracellular trafficking route than the fluorescent liposomal tracer dye^51^, which remains within the membrane while the cargo is released into the cytoplasm. Thus, to verify true cargo delivery, RSV uptake had to be quantified separately from the membrane-bound fluorescent dye examined above. RSV capability to reduce oxidative stress in hyperglycemic hCMEC cells was tested by measuring the mitochondrial membrane potential recovery using ApoE-FL carrier particles with and without RSV payload. First, mitochondrial oxidative stress in hCMEC/D3 cells was induced by incubation with highly concentrated glucose solutions (50 mM and 150 mM). Cells were post-treated with ApoE-FL or ApoE-FL/RSV and mitochondrial depolarization was monitored using JC-1 ratiometric indicator (Extended Data Fig. 1G–H). Significant mitochondrial membrane depolarization was induced at a glucose concentration of 150 mM and only ApoE/FL/RSV treatment was able to normalize it. Particles lacking in RSV did not induce any beneficial changes in cellular mitochondrial functions. Our analysis demonstrated that RSV is successfully delivered into the cytoplasm of hCMEC cells via direct liposome docking and subsequent fusion with the plasma membrane, enabling its downstream effects on mitochondrial function and oxidative stress.

### ApoE-FL/RSV and FL/RSV primarily target the cerebrovascular system in the brain

The biodistribution of liposomes is a critical factor determining their efficacy as drug-delivery vehicles. Moreover, understanding the organ-specificity of liposomes provides insight into their targeting mechanism. Therefore, we first investigated the *in vivo* organ specific distribution of FL and ApoE-coated FL, both loaded with RSV and labeled with the fluorescent dye DiD. Building on the insights gained from our *in vitro* studies, we selected the ApoE-FL/RSV formulation with an ApoE protein-to-lipid ratio of 1:5,000 mol/mol and a lipid-to-RSV ratio of 2:0.4 mol/mol. This specific formulation was chosen based on its optimal physicochemical properties and stability, making it well-suited for in vivo evaluation of both targeting efficiency and anti-aging effects. C57BL/6 mice (5 months old, N=5–7/group) were used, with each receiving a retro-orbital injection of 120 µL of DiD-labeled FL/RSV and ApoE-FL/RSV at a high liposome concentration (10 mg/mL) to ensure sufficient fluorescent signal to noise ratio for visualization. A 30-minute post-injection time point was chosen to focus on early vascular fusion and to avoid liposomal clearance or excretion effects. We found that both FL/RSV-DiD and ApoE-FL/RSV-DiD liposomes resulted in measurable fluorescence signals in all organs (Fig. 3A–E) with significantly higher intensity in the brain for ApoE-FL/RSV compared to FL/RSV (**p<0.01 or ***p<0.001, Fig. 3D–E). This indicates that ApoE functionalization enhances brain-targeting efficiency.

**Figure 3.**
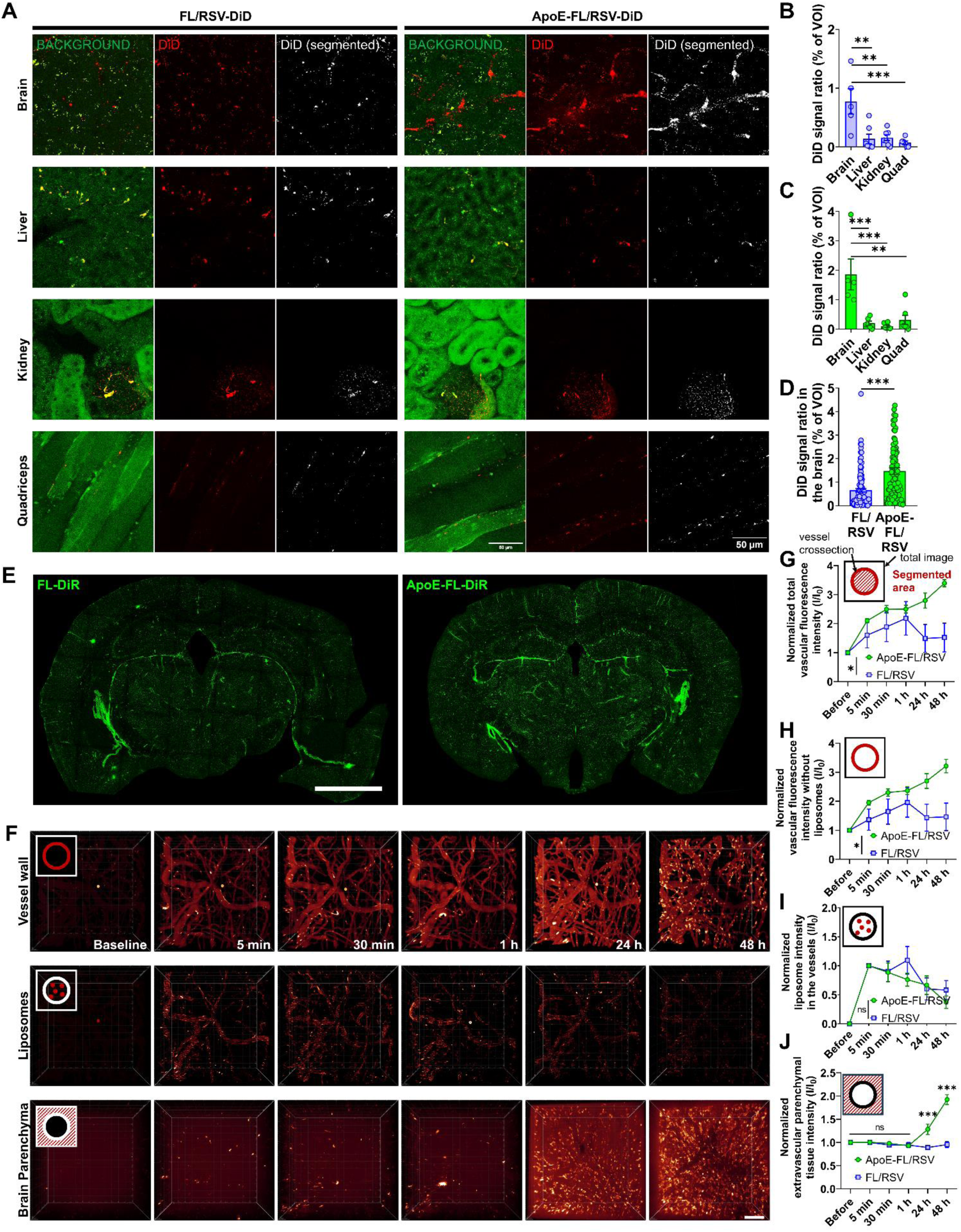
Organ-level biodistribution and brain vascular signal kinetics of DiD/DiR-labeled fusogenic liposome formulations. **A)** Representative images showing the distribution of DiD-derived fluorescence from FL-DiD and ApoE-FL-DiD in sectioned brain, liver, kidney, and quadriceps muscle. These images illustrate organ-level differences in the presence of liposome-associated fluorescent signal. **B–D)** Quantitative analysis of DiD fluorescence in brain, liver, kidney, and quadriceps muscle tissue, derived from voxel-based segmentation. For each section, DiD-positive voxels were identified using an intensity threshold determined from background-corrected control tissue. The percentage of DiD-positive voxels relative to total brain volume was calculated. **E)** Representative whole-coronal brain sections from mice treated with FL-DiR or ApoE-FL-DiR. The ApoE-FL-DiR group exhibits qualitatively stronger and more widespread DiR-associated fluorescence compared with the FL-DiR group. Scale bar: 500 µm. **F)** 3D segmentation of the vasculature in mouse brain with iv injected FL-DiD or ApoE-FL-DiD. Fluorescence signal from the cerebrovasculature has been recorded as a TPM static z-stack at different time points and segmented with IMARIS software in 3D to exclude non-vascular related regions based on 500 kDa FITC-dextran tracer intensity. Scale bar represents 100 μm. **G)** Normalized total vascular fluorescence intensity over time following FL-DiD or ApoE-FL-DiD injection. After 3D vascular segmentation, the cumulative DiD fluorescence within the vascular compartment was normalized first to total segmented vascular volume and then to each mouse’s baseline (pre-injection) signal. **H)** Normalized vascular fluorescence intensity after removing liposome-associated signal. Vessel structures were segmented in 3D, and liposomes (moving and static) were identified and excluded using supervised machine learning based on shape, size, and orientation. The remaining DiD signal represents membrane-integrated fluorescence within endothelial structures. Intensities were normalized using the same procedure as in panel G. **I)** Normalized intravascular liposome intensity over time. Liposomes identified through double 3D segmentation (vascular segmentation followed by liposome-specific machine learning segmentation) were quantified and normalized to their signal immediately after injection (time 0). This captures the clearance kinetics of DiD-labeled liposomes from the vascular space. **J)** Normalized extravascular (parenchymal) DiD fluorescence. After segmenting and removing the entire vascular compartment, the remaining DiD signal in the brain parenchyma was measured and volume-normalized. This represents DiD presence outside the vasculature over time. A marked increase at 24 h is observed, particularly with ApoE-FL-DiD. Across all panels, ApoE-FL-DiD exhibits consistently higher vascular retention than FL-DiD, while parenchymal DiD accumulation at later time points (24 h) is more pronounced with ApoE-functionalized liposomes. Liposome-specific signals show similar clearance profiles for both formulations. Data are presented as mean±SEM (N=7). Statistical significance: *p<0.05, **p<0.01, ***p<0.001 (one-way ANOVA or two-sample t-test).

To determine whether immune-cell sequestration contributes to liposome biodistribution, we examined the interaction of FL and ApoE-FL liposomes with immune cells in the blood and spleen using flow cytometry. In circulating blood cells, we observed a transient fluorescent signal, consistent with brief surface attachment rather than actual uptake, that disappeared within 30 minutes after retro-orbital injection (Extended Data Fig. 4A, C). In the spleen, a modest increase in DiO fluorescence was detected only in the FL-treated group at the 2-hour time point, whereas ApoE-FL liposomes showed no appreciable accumulation (Extended Data Fig. 4B, D). These findings indicate that neither formulation undergoes meaningful immune-cell uptake and that ApoE functionalization minimizes peripheral immune interactions, supporting its enhanced suitability for targeted brain delivery.

To validate endothelial-specific targeting in the brain, we utilized a vascular endothelial cadherin (VECAD)-tdTomato transgenic mouse model, where endothelial cells specifically express the fluorescent tdTomato protein. To ensure signal specificity and rule out potential imaging artifacts, we incorporated additional lipid-based fluorescent tracers, DiR and DiO, into the liposomal formulation. Using two-photon microscopy, we confirmed effective targeting of brain endothelial cells by both FL and ApoE-FL liposomes, as evidenced by their colocalization with tdTomato-labeled endothelial cells in cortical and subcortical vessels (Extended Data Fig. 6). To further evaluate cell-type specificity, we analyzed the colocalization of the DiD fluorescent signal in the brain with markers for astrocytes (glial fibrillary acidic protein; GFAP), microglia (ionized calcium-binding adaptor molecule 1; IBA1), and neurons (Neuronal nuclei; NeuN), and found no substantial overlap (Extended Data Fig. 7A), indicating no off-target accumulation in these cell types.

As expected from RSV’s known systemic vascular effects, both FL-RSV and ApoE-FL/RSV improved aortic relaxation (Extended Data Fig. 8A–C), confirming effective peripheral delivery of the therapeutic payload. Importantly, beyond the systemic actions, the engineered liposomes demonstrated a distinct and much more selective impact within the brain microvasculature. Fusogenic liposomes, whether coated with a spontaneous or engineered ApoE protein corona, showed robust endothelial-specific accumulation in the brain. The fluorescent signal from FL was consistently lower than that of ApoE-FL, suggesting less efficient targeting. In contrast, ApoE-functionalized liposomes showed significantly enhanced adhesion and specificity, likely driven by ApoE–LRP1 receptor-mediated interactions. This targeted enhancement highlights the advantage of incorporating ApoE into the protein corona, effectively optimizing the liposomes for more efficient and selective delivery to the brain.

### *In vivo* fusion dynamics of ApoE-FL/RSV and FL/RSV liposomes along the cerebral arteriovenous axis

To investigate the *in vivo* fusion dynamics of ApoE-FL/RSV-DiD and FL/RSV-DiD liposomes along the cerebral microvascular arteriovenous axis, we employed intravital two-photon microscopy to track their movement and accumulation in the cerebral vasculature and brain parenchyma. This approach allowed us to evaluate the dynamics of liposome fusion by quantifying the localization and intensity of DiD fluorescence signal over time (Fig. 3F–J). For quantification, DiD intensity changes were monitored for 48 hours post-administration (0, 0.5, 1, 24, and 48 h), and 3D segmentation analysis was performed to assess signal intensity in the vasculature (Fig. 3G) (moving liposomes vs. static labeling) and extravascular brain parenchymal tissue (Fig. 3J). To ensure signal specificity and exclude imaging artifacts, additional fluorophores (DiR and DiO) were used for validation in separate experiments.

DiD intensity of moving liposomes in the vascular lumen peaked immediately after injection, and gradually decreased over time, indicating the reduction of circulating liposomes (Fig. 3I). At the same time the vascular wall intensity (total vascular DiD intensity minus the luminal intensity) increased significantly in both FL and ApoE-FL injected mice (Fig. 3H), reflecting fusion with endothelial cell membrane. This signal accumulation was substantially more pronounced in the ApoE-FL treated group, demonstrating superior targeting and fusion capability of the ApoE protein corona-functionalized liposomes. Fluorescent intensity changes in the extravascular brain parenchyma (total intensity minus vascular intensity) only showed a notable increase 24 hours after injection (Fig. 3J). This delayed appearance of fluorescent signal in the brain parenchyma suggests that the underlying mechanism is not endothelial transcytosis, which typically occurs within 10 to 30 minutes in the BBB^54, 55^. Instead, this prolonged timeline indicates involvement of a much slower membrane recycling or lipid redistribution process. Since DiD is a lipophilic membrane dye that stably incorporates into the lipid bilayer, its delayed detection beyond the endothelium implies gradual lateral diffusion or membrane turnover rather than vesicle-mediated transport. These findings support the hypothesis that fusogenic liposomes merge with the endothelial luminal plasma membrane, delivering their lipid components and the DiD tracer directly into the membrane. Following fusion, DiD diffuses laterally within the endothelial membrane and can move independently of the bulk liposomal lipids, gradually redistributing toward the abluminal surface or entering the parenchyma through slower membrane–trafficking processes.

Vessel microanatomy and fluid biomechanics differ between arterioles, capillaries, and venules. This principal feature of the brain is often overlooked in drug delivery studies. The limited data available for cationic liposomes imply that they are taken up only by post-capillary venules^56^. Here, to address these questions, we used two-photon *in vivo* imaging to examine in the intact brain how distinct types of cerebral vessels handle FL/RSV and ApoE-FL/RSV in real-time (Fig 4.). We show that both ApoE-FL/RSV and FL/RSV liposomes fuse primarily with endothelial cells in capillaries and post-capillary venules, vascular segments defined by low-flow, low-pressure conditions that favor endothelial trafficking and local nanoparticle fusion. (Fig. 4). The ability of fusogenic liposomes to be taken up by both capillaries and post-capillary venules, unlike traditional cationic liposomes, which are primarily restricted to uptake at post-capillary venules^56^, significantly expands the potential surface area for brain delivery. Capillaries constitute the vast majority of the cerebrovascular network, with a density of approximately 500–600 mm/mm³ ^57^ and an estimated total length of around 150 meters in the mouse brain alone^58^. In contrast, post-capillary venules represent a much smaller fraction of the vascular bed, both in terms of density and total length. Therefore, the dual uptake capacity of fusogenic liposomes enables access to a substantially larger vascular interface, potentially enhancing efficiency, uniformity, and distribution of therapeutic delivery across the brain parenchyma. This broader engagement with the cerebrovascular system is particularly advantageous for achieving widespread and effective targeting in neurovascular and neurodegenerative conditions.

**Figure 4.**
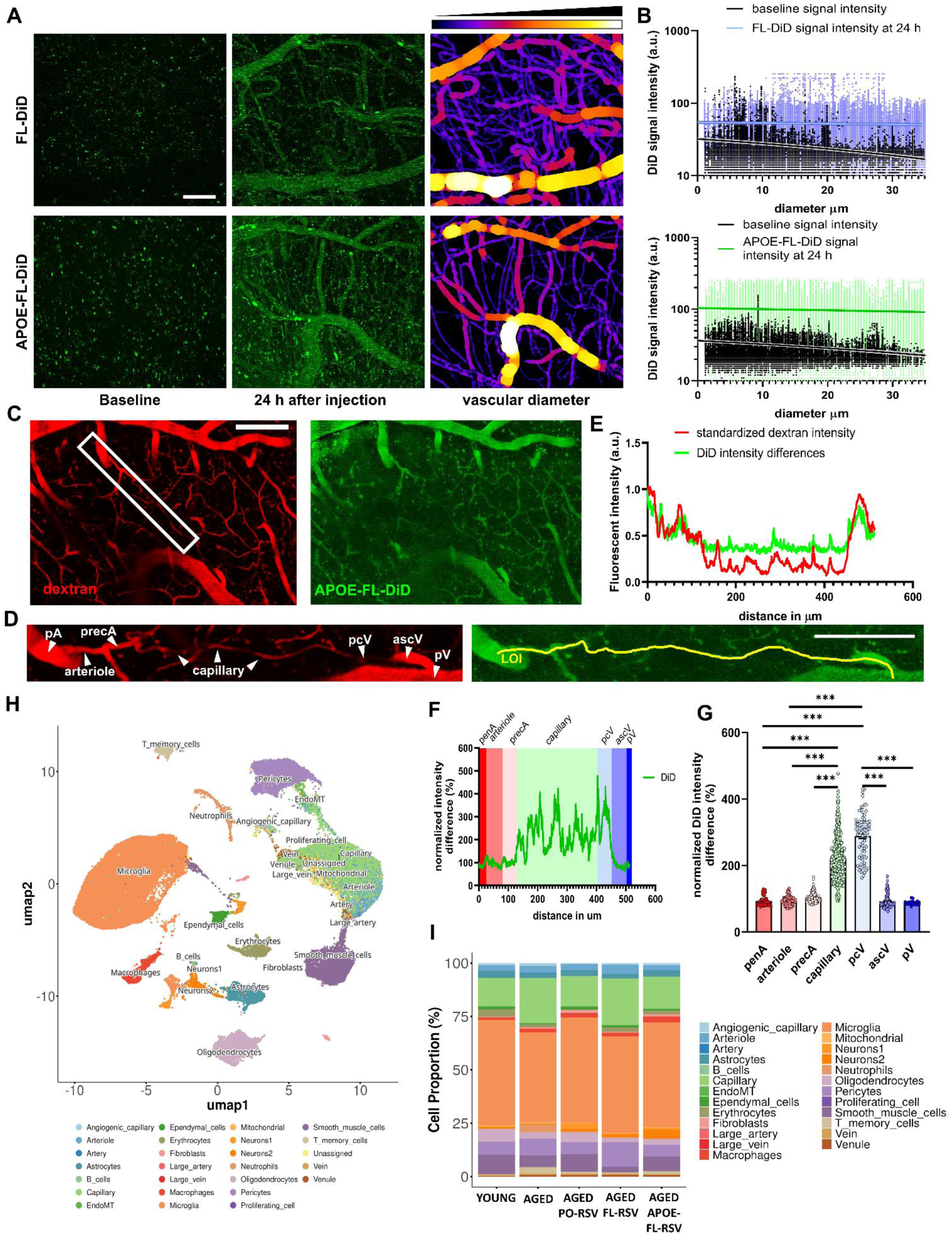
Distribution and vascular localization of ApoE-FL-DiD in the cerebromicrovasculature. **A)** Representative two-photon microscopy images showing the distribution of FL-DiD and ApoE-FL-DiD nanoparticles in cerebral microvessels at baseline (before injection) and 24 hours post-injection together with vascular diameter mapping. Scale bar: 50 µm. **B)** Quantitative analysis of DiD fluorescence intensity as a function of vessel diameter at baseline (black line) compared to 24 hours after injection of FL-DiD and ApoE-FL-DiD. Signal intensity values are presented as scatter plots with fitted trend lines. **C)** Representative high-magnification images showing colocalization of vascular dextran tracer (red) and ApoE-FL-DiD nanoparticles (green) to assess the vascular zonation of fusion events within a region of interest (white rectangle) in the cerebral vasculature. **D)** Representation of the magnified view of the analyzed vessel segment. Penetrating arteriole (pA), arteriole, pre-capillary arteriole (precA), capillary, post-capillary venule (pcV), ascending venule (ascV), and penetrating venule (pV) are clearly identifiable as a line of interest (LOI). **E)** Graph showing standardized dextran intensity (red line) versus ApoE-FL-DiD intensity (green line) measured along the vessel segments. **F)** Normalized DiD intensity along the analyzed vessel segments, color-coded by vessel zonation. **G)** Statistical analysis comparing normalized DiD intensity among different vessel zones, demonstrating significant preferential localization of ApoE-FL-DiD nanoparticles within the capillary and a pcV. Data shown as mean±SEM; statistical significance determined by ANOVA with Tukey’s multiple comparison test. ***p<0.001. **H)** Single-cell transcriptomic mapping of cerebrovascular and neural cell populations. UMAP visualization of single-cell RNA-seq data showing distinct transcriptional clusters across major brain cell types, including endothelial subtypes (arteriole, capillary, venule), perivascular cells, microglia, astrocytes, neurons, oligodendrocytes, fibroblasts, and immune cells. Each cluster is color-coded according to its annotated identity, illustrating the cellular heterogeneity captured in the dataset. **I)** Quantitative comparison of cell-type composition across experimental groups. Stacked bar plots showing the proportional distribution of single-cell transcriptomic clusters in young, aged, and aged + treatment groups (PO-RSV, FL-RSV, APOE-FL-DiD). Aging is associated with reduced endothelial and pericyte fractions and an expansion of glial and immune populations. APOE-FL-DiD treatment partially restores endothelial representation and reduces microglial prevalence, suggesting improved vascular–glial homeostasis. The partial normalization of cellular composition in treated groups suggests attenuation of neuroinflammatory remodeling.

Single-cell RNA-seq analysis (Fig. 4.H–I) further supported these findings. Clustering identified distinct endothelial, pericyte, and glial populations (Extended Data Figure 5), and quantitative comparison across groups showed that reduced endothelial and pericyte fractions while expanding inflammatory microglia, consistent with neurovascular aging. Treatment with ApoE-FL-RSV partially reversed these changes, restoring endothelial representation, reducing microglial abundance, and re-establishing vascular–glial balance toward the young phenotype. Together, the imaging and transcriptomic data indicate that capillary and venular endothelial cells are the principal sites of liposome fusion, and that their engagement by ApoE-FL-RSV not only broadens vascular access but also may mitigate age-related neuroinflammatory remodeling, supporting improved cerebrovascular homeostasis.

### ApoE-FL/RSV and FL/RSV rejuvenate structural and functional cerebrovascular impairment in aged mice

To maximize the potential of ApoE-FL as a delivery system, it is crucial to assess potential changes in LRP1 expression within the aged brain vasculature, as an age-related decline in LRP1, such as that reported in Alzheimer’s disease^59^, could impact drug delivery efficiency. Our analysis demonstrated that during mouse brain aging, LRP1 expression remained relatively stable at both mRNA (Extended Data Fig. 2H) and protein level (Extended Data Fig. 2I). Reanalysis of publicly available human brain scRNA-seq datasets further supported this finding, showing that brain endothelial LRP1 expression is preserved with age and exhibits only a slight decrease by age 90 (Extended Data Fig. 2J). This consistent expression of LRP1 across aging suggests that it is unlikely to significantly influence the binding capacity or delivery efficiency of ApoE-FL in the aged brain.

#### 1. ApoE-FL/RSV and FL/RSV improve blood-brain barrier integrity in aged mice

While RSV has shown promise in restoring BBB integrity^60^, its poor bioavailability limits its therapeutic efficacy when administered orally^61^. We evaluated the potential of RSV delivered via advanced fusogenic liposomal systems, FL-RSV and ApoE-FL-RSV, to restore BBB integrity in aged mice, in comparison to conventional oral RSV administration. Notably, we used an oral RSV dose that was approximately 2000 times higher than the liposomal dose (200 mg/kg body weight per day orally vs. 0.117 mg/kg per day delivered via a 2 mg/mL liposome-RSV complex), allowing us to assess the efficiency of targeted delivery relative to high-dose systemic exposure. To assess BBB integrity, we employed a well-established intravital two-photon microscopy technique^62^, which enables high-resolution, real-time visualization of BBB permeability *in vivo*. Fluorescein isothiocyanate (FITC)-dextran tracers of varying molecular weights (0.3 kDa to 40 kDa) were administered via retro-orbital injection, and their extravasation into the brain parenchyma was monitored to evaluate size-selective barrier permeability under both physiological and pathological conditions. For experimental groups, 6 months old mice served as young controls, while untreated 24 months old mice were used as aged controls.

Remarkably, liposomal treatment led to a significant reduction in vascular tracer extravasation and improved BBB integrity as early as five days after the start of daily administration, lowering BBB permeability in aged animals to levels comparable to those of young controls (Fig. 5A–B). To assess the long-term effects, animals were treated for one month, with oral RSV included as a chronic treatment control. While oral RSV resulted only in modest improvements in BBB permeability both FL/RSV and ApoE-FL/RSV treatments proved substantially more effective (Fig. 5C). Notably, ApoE-FL/RSV treatment showed a particularly significant effect even in the smallest tracer size range, further supporting the advantage of ApoE-mediated targeting in enhancing therapeutic outcomes (Fig. 5A–C).

**Figure 5.**
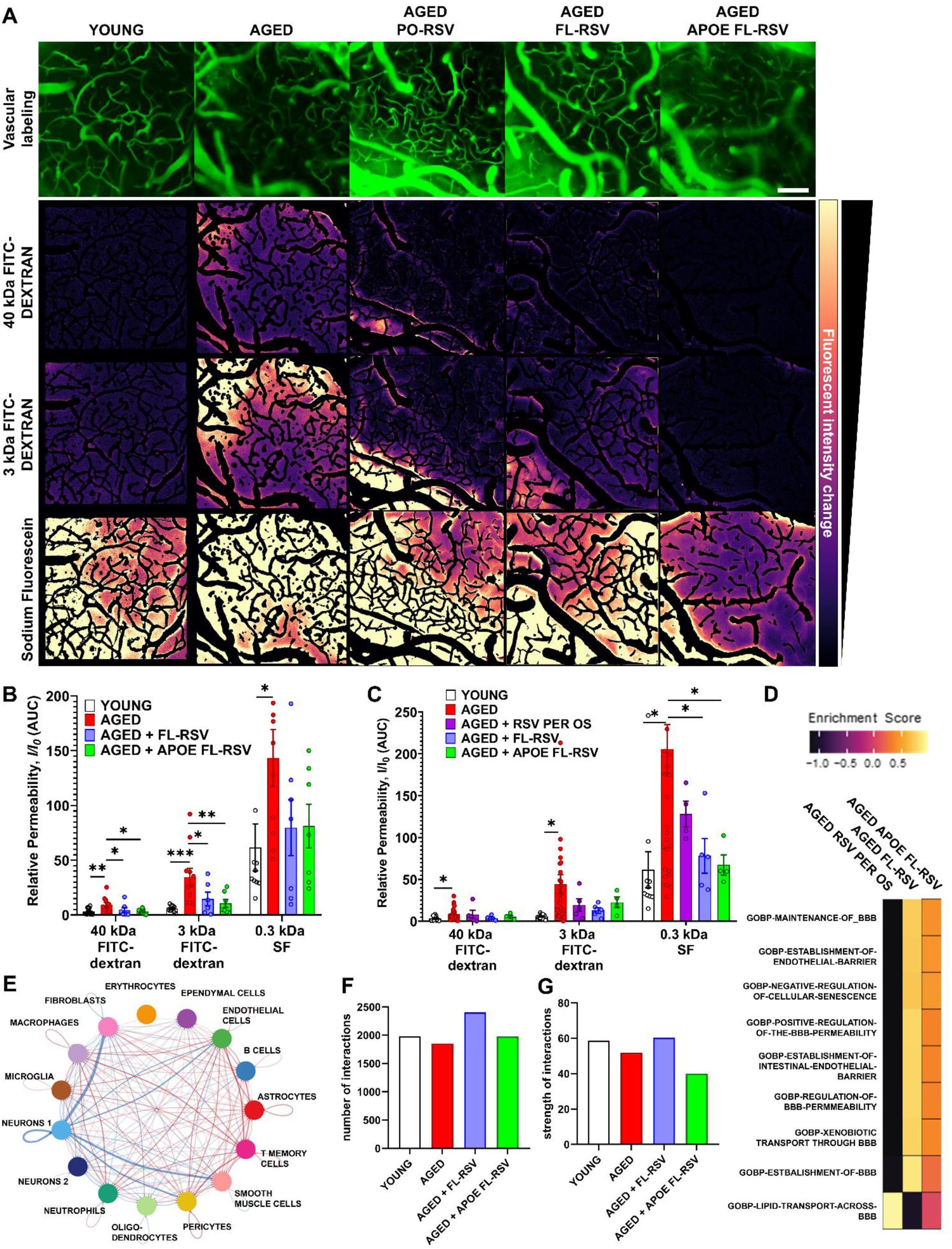
FL-RSV and ApoE-FL-RSV significantly improve BBB integrity in aged mouse brains. Two-photon microscopy-based measurement of microvascular permeability to fluorescent tracers in brains of young control and aged mice that received the vehicle, RSV *per os* (for 1 month), FL-RSV or ApoE-FL-RSV. **A)** Representative intensity maps of the mouse brain during microvascular permeability measurement. Treatment with FL-RSV and ApoE-FL-RSV notably reduces extravascular tracer intensities in maximum projection images, indicating a decrease in vascular permeability. The scale bar represents 100 μm. B-C) Quantification of relative permeability changes in aged mice treated with FL-RSV, and ApoE-FL-RSV for 4 days **(B)** and 1 month **(C)**. In aged mice, RSV treatment delivered with FL or APOE-FL effectively improved BBB permeability for all tracers tested both after 4 days and 1-month treatment, while RSV *per os* only partially improved the barrier permeability for 3 and 0.3 kDa tracers but not for 40 kDa. **D)** Gene ontology enrichment analysis highlights those protective pathways related to BBB maintenance, endothelial barrier formation, and negative regulation of cellular senescence are most enriched in ApoE-FL/RSV and FL/RSV groups. In contrast, oral RSV shows comparatively weaker enrichment of these protective processes and higher enrichment in pathways associated with positive regulation of BBB permeability, suggesting a less stable barrier. **E–G)** Downstream network analysis of single-cell RNA sequencing (scRNA-seq) demonstrates increased cellular connectivity among key BBB-associated populations like endothelial cells, pericytes, and astrocytes in ApoE-FL/RSV and FL/RSV-treated brains, indicating enhanced intercellular communication and functional integration. Number of cell-cell interaction and the strength of these intercellular connections reveal that endothelial cells from ApoE-FL/RSV and FL/RSV-treated groups exhibit higher engagement, supporting improved BBB structural cohesion. These findings support that liposomal RSV treatment, particularly with the ApoE-enriched formulations, enhances BBB function in aging, likely via improved cellular communication and endothelial barrier reinforcement. Data are representing mean±SEM, N>7 for all groups. *p<0.05, **p<0.01, ***p<0.001 with Kruskal-Wallis test.

To further characterize the molecular changes in the BBB we performed scRNA-seq analysis by using cell network analysis and gene ontology (GO) enrichment (Fig. 5D–E). Network analysis reveals that ApoE-FL/RSV and FL/RSV exhibit increased connectivity among key BBB-associated cells, such as endothelial cells, pericytes, and astrocytes, indicating enhanced cellular crosstalk and functional integration. Quantitative analysis of both the number and strength of intercellular interactions further supported this observation demonstrated elevated interaction metrics in the ApoE-FL/RSV and FL/RSV treatment groups compared to nontreated aged controls (Fig. 5 F–G). The GO enrichment analysis indicates that pathways related to BBB maintenance, endothelial barrier establishment, and negative regulation of cellular senescence are significantly enriched in ApoE-FL/RSV and FL/RSV-treated mice, whereas oral RSV treatment exhibits weaker enrichment of these protective pathways. Together, these findings suggest that liposomal RSV delivery, particularly due to ApoE in the protein corona, enhances BBB function in aging more effectively than oral RSV, likely by improving cellular interactions and reinforcing endothelial barrier integrity.

#### 2. ApoE-FL/RSV and FL/RSV restore vascular density and enhance neurovascular coupling responses in aged mice

Age-related cognitive decline is strongly linked to decrease in CBF either due to structural and/or functional alterations. Structural changes include microvascular rarefaction, the progressive loss of microvascular density in the brain^32^. This reduction in vascular density leads to impaired basal CBF, limiting the delivery of oxygen and nutrients to neurons, and contributing to neuronal dysfunction^32^. Compounding this is the functional deterioration of the cerebromicrovascular system including the decrease in NVC, the mechanism by which blood flow dynamically adjusts to meet the metabolic demands of active neurons. Impaired NVC exacerbates neuronal dysfunction, underscoring the critical importance of maintaining vascular integrity and NVC in aging brains to prevent or mitigate cognitive deficits^32^. Microvascular density was evaluated using ICONEUS Ultrasound Localization Microscopy (ULM), a novel intravital imaging technique, utilizing intravenously injected microbubbles, which scatter ultrasound waves as they traverse the blood vessels, generating high-resolution vascular images with a spatial resolution of 2 µm/pixel (Fig. 6A). This technique enables real-time intravital imaging of not only cortical microvasculature but also subcortical regions, including the hippocampus, allowing for a comprehensive assessment of cerebrovascular architecture in deeper brain structures. Vascular density in the cortical (Fig. 6B) and hippocampal (Fig. 6C) regions was quantified by analyzing microbubble intensity maps using ImageJ, with results expressed as percentage signal changes per volume of interest (VOI).

**Figure 6.**
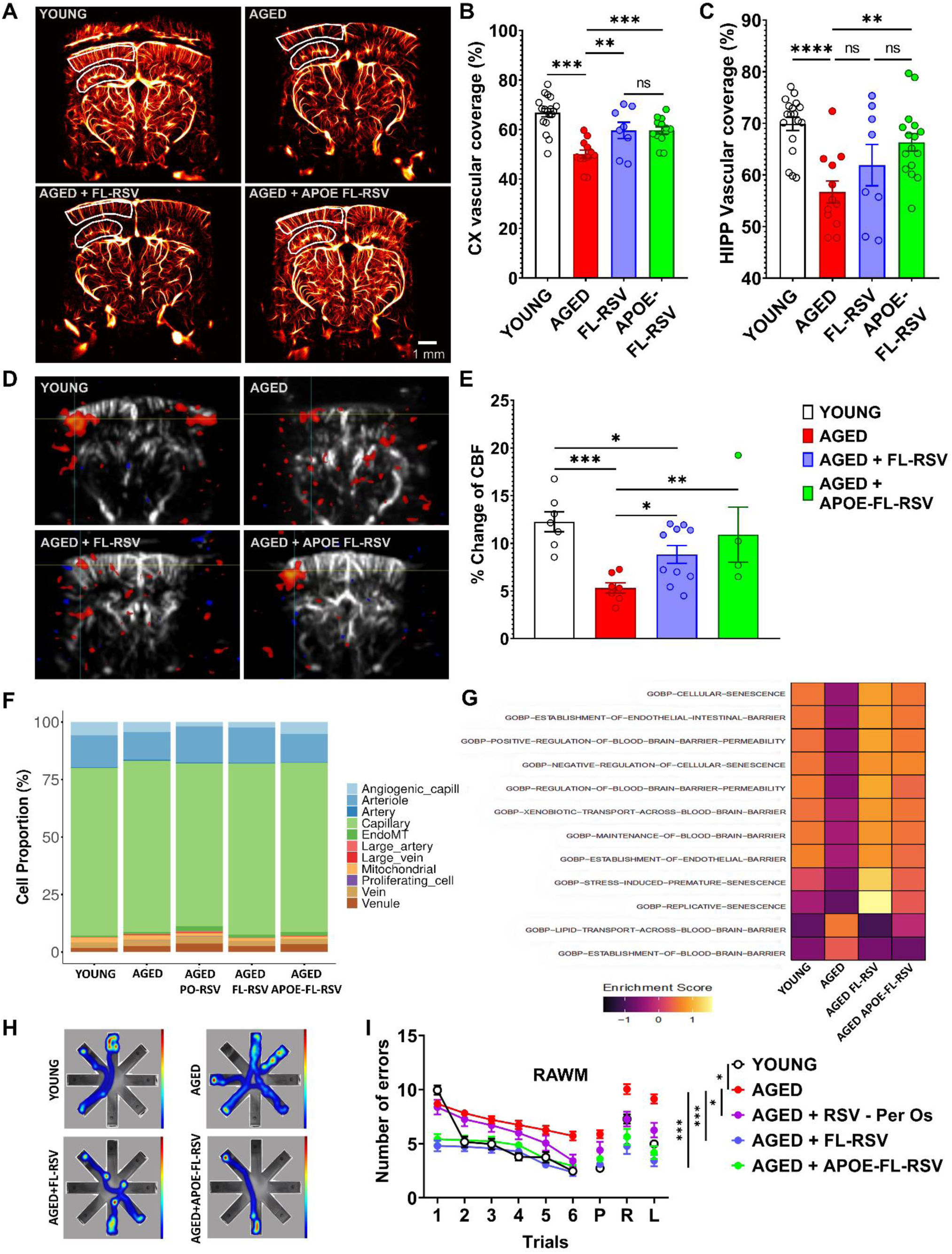
FL-RSV and ApoE-FL-RSV partially restored neurovascular coupling responses, improved vascular density and enhanced learning and memory capabilities in aged mice. **A)** Representative images of the cerebral microvasculature recorded by functional ultrasound imaging resulting in the presented ultrasound localization microscopy images. Vascular density has been measured in the cortical (CX) and hippocampal (HIPP) areas of the mouse brain. **B–C)** Quantification of the cortical and hippocampal cerebral vascular density in young, aged, aged + FL-RSV and aged + ApoE-FL-RSV treated animals. **D–E)** Measurement of neurovascular coupling responses in the young, aged, aged + FL-RSV and aged + ApoE-FL-RSV treated mice. Note the significant differences between aged, aged + FL-RSV and aged + ApoE-FL-RSV. Data is shown in mean±SEM. *p<0.05, **p<0.01, ***p<0.001 with ANOVA, N≥8 for all groups. **F)** Single-cell RNA-seq analysis reveals restoration of endothelial heterogeneity with treatment. Stacked bar plots showing the proportional abundance of vascular cell populations across experimental groups. Aging markedly reduces angiogenic and proliferating endothelial subtypes, whereas FL-RSV and particularly APOE-FL-RSV treatment partially restores endothelial diversity and the presence of regenerative cell states, suggesting improved vascular maintenance capacity. **G)** Gene ontology enrichment indicates reduced endothelial senescence and improved barrier maintenance after APOE-FL-RSV treatment. Heatmap of endothelial GO term enrichment scores demonstrates that aging increased pathways associated with cellular senescence, endothelial stress, and barrier dysfunction, while APOE-FL-RSV shifted expression toward gene sets linked to blood–brain barrier integrity, lipid transport, and endothelial homeostasis. **H-I)** Measurement of learning and memory functions of the young, aged, aged + FL-RSV, aged + ApoE-FL-RSV treated animals and aged animals that have RSV *per os*. Young control and aged BL6 mice were subjected to vehicle, RSV *per os*, FL-RSV and ApoE-FL-RSV treatments and assessed for their spatial learning and memory using the Radial Arm Water Maze (RAWM). During the learning phase (days 2 to 6) and on probe (P), retrieval (R) and relearn (L) days aged mice displayed higher combined error rates compared to young mice. Treatment with FL-RSV and ApoE-FL-RSV significantly enhanced learning performance in aged mice relative to their untreated counterparts. The combined error rate was computed by adding one error for each incorrect arm entry plus an error for every 15 seconds of inactivity. Data are presented as mean±SEM (N=10–15 per group). Statistical significance indicated by *p<0.05, **p<0.01, ***p<0.001 using Repeated Measure ANOVA, demonstrating the beneficial effects of FL-RSV and ApoE-FL-RSV treatments on enhancing cognitive functions in aged mice, particularly in spatial learning, but less so in tasks requiring cognitive flexibility.

Our findings demonstrated that both FL/RSV and ApoE-FL/RSV treatments significantly increased cortical vascular density compared to untreated aged mice (p<0.01 and p<0.001, respectively; Fig. 6B), indicating that fusogenic liposome-mediated resveratrol delivery effectively restores cortical microvascular networks. However, regional differences emerged when assessing deeper brain structures. In the hippocampus, only ApoE-FL/RSV treatment enhances significantly the vascular density relative to untreated aged mice (p<0.01, Fig. 6C), while FL/RSV alone failed to restore hippocampal vascularization. This suggests that the engineered ApoE protein corona improves brain endothelial targeting, membrane fusion, and vascular retention of the liposomes, especially in subcortical regions that are otherwise poorly reached by FLs with spontaneous coronas. These results underscore the importance of ApoE functionalization for delivering therapeutic payloads to deeper brain structures, such as the hippocampus, where vascular decline contributes to age-related cognitive impairment.

Neurovascular coupling was evaluated using the same ICONEUS system to measure CBF changes in the somatosensory cortex under baseline conditions and following a 30-second whisker stimulation, which is a well-established model of neuronal activation. CBF responses were quantified as percentage changes from baseline, and corresponding CBF maps (Fig. 6D) illustrated the spatial distribution of vascular responses. Aged mice showed significantly impaired NVC compared to young controls (p<0.001). Treatment with both FL/RSV and ApoE-FL/RSV partially restored NVC in aged animals (p<0.05 vs. aged controls), with ApoE-FL/RSV achieving NVC responses nearly indistinguishable from those of young mice, indicating superior functional recovery (Fig. 6E). Together, the vascular density measurements (Fig. 6A–C) and NVC assessments (Fig. 6D–E) demonstrate that ApoE-FL/RSV treatment not only restores the structural microvascular network but also re-establishes functional perfusion responsiveness.

Consistent with these imaging results, our single-cell RNA-seq data provide strong evidence that ApoE-FL/RSV treatment enhances cerebrovascular density by increasing the proportion of angiogenic and proliferating endothelial cells (Fig. 6F). Compared with both aged and oral RSV groups, only the ApoE-functionalized formulation expanded markedly the angiogenic capillary endothelial subpopulation, indicating activation of endothelial renewal and vascular remodeling programs critical for reversing age-related microvascular decline. The FL/RSV treated group showed a modest shift in endothelial composition, with no significant increase in angiogenic subsets. Additionally, ApoE-FL/RSV treatment increased the proportion of capillary endothelial cells, suggesting improved microvascular maintenance and stability across the vascular network. These effects are consistent with enhanced endothelial targeting and fusion efficiency mediated by the ApoE corona.

Pathway enrichment analysis of endothelial transcriptomes further revealed that aging was associated with increased activation of pathways related to cellular senescence, oxidative stress, and BBB dysfunction (Fig. 6G). In contrast, both FL/RSV and ApoE-FL/RSV shifted endothelial gene expression toward a more youthful profile, characterized by upregulation of pathways involved in angiogenesis (Extended Data Figure 7B), endothelial migration, lipid transport, xenobiotic clearance, and barrier maintenance, while suppressing senescence-related signaling. These transcriptional shifts, together with the expanded angiogenic and capillary endothelial compartments, suggest that fusogenic liposomal RSV delivery not only promotes structural vascular regeneration but also rejuvenates endothelial function. The transcriptional and structural data together demonstrate that ApoE-FL/RSV exerts coordinated effects, restoring basal perfusion capacity, improving neurovascular reactivity, and rejuvenating endothelial function, thereby reinstating cerebrovascular health in aged mice.

### ApoE-FL/RSV and FL/RSV ameliorates age-related learning and working memory deficits in aged animals

Aging is associated with progressive declines in cognitive functions, including memory and spatial learning, driven by both neuronal and vascular changes in the brain^63^. RSV has shown potential for mitigating age-related cognitive decline^64^, although its efficacy is limited due to poor bioavailability. Cognitive functions such as spatial learning and short- and long-term memory, were tested by assessing performance in the radial arm water maze (RAWM) (Fig. 6H), following our published protocols^65^. The RAWM assesses spatial learning, short-term memory, and long-term memory in rodents. Spatial learning is measured by reduced latency and errors across trials, while short-term memory is evaluated by performance improvements within a session. Long-term memory is tested in probe trials conducted 24 hours or later, assessing memory retention. The results (Fig. 6I) indicate a significant impairment in spatial learning and memory in aged control mice compared to young controls. During the learning phase (trials 2–6), aged mice made substantially more errors than young mice (p<0.001), highlighting the cognitive deficits associated with aging. This impairment persisted during the probe (P), retrieval (R), and relearn (L) tests, in which aged controls continued to exhibit significantly higher error rates compared to young controls (p<0.001), confirming the negative impact of aging on both memory consolidation and recall.

Treatment with oral RSV significantly improved performance compared to untreated aged mice, as evidenced by fewer errors throughout the trials. However, RSV-treated mice still performed worse than young controls, likely due to the limited bioavailability of oral RSV, which may have restricted its therapeutic efficacy. In contrast, both FL/RSV and ApoE-FL/RSV treatments resulted in a substantial reduction in error rates across all phases of the task, with performance levels becoming statistically indistinguishable from young controls (p<0.05 to p<0.001 vs. aged controls).

Decreases in swimming speed due to age-related skeletal muscle dysfunction can confound studies assessing cognition using the RAWM. We evaluated swimming speed and found that it slightly decreased in aged mice vs. young but it was not different between aged mice and aged mice treated with FL-RSV or ApoE-FL-RSV (Extended Data Fig. 8D). This confirms that the observed cognitive improvements are not influenced by physical performance. These findings highlight the superior efficacy of targeted liposomal RSV delivery in reversing age-related cognitive decline.

## Discussion

Age-related cognitive impairment is becoming increasingly prevalent as the aging population grows^66–68^, and has been linked to cerebrovascular decline^69, 70^. While direct intervention at the neuronal level remains challenging, the vascular component of cognitive decline offers a promising therapeutic target^4, 9, 10, 13, 65, 71–75^. By improving age-related cerebrovascular function and structure, there is potential to enhance cognition. Although several naturally occurring compounds exhibit vasoprotective effects, such as resveratrol, their low bioavailability and stability limit their use in targeted cerebrovascular therapy. To address this, we developed a novel delivery system using fusogenic liposome formulations. By leveraging the properties of an *in vivo* spontaneously forming protein corona enriched in apolipoproteins, we created an ApoE-FL/RSV delivery system, demonstrating its fundamental impact on cerebrovascular aging and cognitive health.

We have previously characterized the beneficial effects of FL/RSV both *in vitro* and *in vivo*^40, 49^. We demonstrated that its direct fusion with endothelial cell membranes achieves intracellular RSV concentrations *in vitro* five times higher than those delivered by conventional cationic liposomes^49^. *In vivo*, we showed that RSV delivered via FL can positively impact the aging brain by improving neurovascular coupling^40^. Building on these findings, we aimed to comprehensively characterize the cerebrovascular effects of membrane fusion-based delivery systems and finetune their brain targeting potential by leveraging their natural propensity to accumulate a passive protein corona enriched with apolipoproteins. Using this approach, we functionalized FL with full-length ApoE protein to enhance their brain-targeting efficiency and therapeutic potential.

Protein corona formation can exert both beneficial and detrimental effects on nanoparticle-based delivery systems. When the corona contains proteins that engage receptor-mediated pathways, it can improve targeting by promoting selective uptake into specific cell types or organs. However, the corona can also reduce targeting efficiency by redirecting nanoparticles to off-target tissues or increasing their routing to lysosomes, which ultimately lowers therapeutic efficacy^52, 76–80^. To explore this, we analyzed the spontaneously forming protein corona around FL liposomes in both human and mouse serum and found that apolipoproteins were significantly enriched compared to other serum proteins. Notably, ApoE, a key apolipoprotein involved in lipid transport across the BBB, was highly abundant in the protein corona. ApoE plays a crucial role in maintaining brain lipid homeostasis, particularly in the transport of lipids to neurons via receptor-mediated endocytosis^81^.

It primarily interacts with low-density lipoprotein receptors, including LDLR, LRP1, VDLR, APOER2, which are abundantly expressed on brain endothelial cells and neurons^82,83^. Through these interactions, ApoE facilitates the delivery and clearance of lipoprotein particles, supports synaptic remodeling, and regulates neuroinflammation^84^, all of which are essential for cognitive function and neuronal health^85, 86^. Given its fundamental role in lipid transport and receptor-mediated endocytosis across the BBB^87^, ApoE-functionalized nanoparticles and liposomes have been shown to improve CNS drug targeting by facilitating receptor-mediated uptake across brain endothelial cells^44, 88, 89^. Our findings reveal that ApoE is highly enriched in the spontaneously formed protein corona of FL, and this enrichment plays a beneficial role in brain-targeted delivery. Rather than impairing FL function or increasing lysosomal degradation, the presence of ApoE in the protein corona facilitates the selective targeting of FL to brain endothelial cells without compromising their fusogenic capacity or facilitating their immune cell uptake and clearance (Extended Data Fig. 6). This mechanism was successfully harnessed to enhance RSV delivery to the brain, supporting the development of ApoE-FL as a targeted therapeutic strategy for cerebrovascular rejuvenation. Furthermore, the protein corona-mediated targeting of brain microvascular structures includes both the entire capillary system and postcapillary venules, enhancing the effectively targeted vascular surface area. Thus, the newly engineered ApoE-FL system combines the natural brain-targeting ability of ApoE with the unique properties of fusogenic liposomes, including direct membrane fusion, efficient intracellular cargo delivery, and minimized lysosomal trapping, offering a powerful platform for age-related neurovascular interventions.

To maximize the potential of ApoE-FL as a delivery system, it is crucial to assess potential changes in LRP1 expression within the aged brain vasculature, as LRP1 is the most prominent ApoE receptor in the brain endothelial cell surface^87^. Our findings demonstrate that LRP1 expression remains stable in the aged mouse brain (Extended Data Fig. 2G–I), suggesting that it is unlikely to significantly affect the binding efficiency and delivery potential of ApoE-FL in age-related cerebrovascular dysfunction. While human studies indicate that LRP1 expression declines in Alzheimer’s disease brain, impacting the brain’s ability to clear amyloid-beta peptides^90^, our analysis of publicly available human scRNA-seq datasets shows that endothelial specific LRP1 expression is maintained in the aging brain (Extended Data Fig. 2J), suggesting that in age-related cognitive decline endothelial ApoE-FL-mediated delivery may not be inherently compromised. This discrepancy may be due to the fact that previous studies assessed LRP1 expression in whole brain tissue extracts, which includes neurons and astrocytes, also known to highly express LRP1, rather than using a cell type-specific approach^91^. As a result, lack of changes in endothelial LRP1 expression may have been obscured by the overall cellular heterogeneity. Currently, the therapeutic effect of RSV appears to be fully maximized with both FL and ApoE-FL formulations in the cortex, achieving near-complete cerebrovascular rejuvenation. Therefore, even in the context of Alzheimer’s disease, where endothelial LRP1 expression may be reduced and ApoE-mediated targeting potentially impaired, ApoE-FL is still expected to deliver significant therapeutic benefit. In such cases, fine-tuning the RSV concentration within the ApoE-FL formulation could help maintain therapeutic efficacy, ensuring that the benefits of RSV are preserved even under conditions of compromised receptor availability.

Although ApoE was selected for CNS targeting through LDLR/LRP-mediated docking and fusion with endothelial cell membrane, similar strategies could be adapted in the future for other organs by exploiting apolipoproteins with tissue-specific receptor tropisms. For example, ApoA-I or ApoA-II could guide liposomal fusion to SR-B1-rich hepatic and pulmonary endothelium^92^ or atherosclerotic lesions^93^, ApoB fragments could facilitate hepatocyte-directed uptake via LDLR^92, 94^, and ApoM could enable selective renal or systemic vascular endothelial targeting through S1P-associated pathways^95^. Such apolipoprotein diversification could broaden the therapeutic potential of fusogenic liposomes beyond the brain.

The ability of fusogenic liposomes to be taken up by both capillaries and post-capillary venules, unlike traditional cationic liposomes, which are primarily restricted to uptake at post-capillary venules^56^, significantly expands the potential surface area for brain delivery. Previous studies have shown that various nanocarriers, including polymeric nanoparticles and antibody-conjugated nanoparticles, primarily undergo transcytosis-mediated brain entry at post-capillary venules, with negligible uptake in capillaries^56, 96^. For example, transferrin receptor-targeted liposomes have been observed to accumulate in the endothelium of capillaries and venules, but actual transcytosis into the brain parenchyma predominantly occurs at post-capillary venules^56^. The ability of fusogenic liposomes to be taken up by both capillaries and post-capillary venules represents a significant advantage over these other nanocarriers, potentially enhancing the efficiency and distribution of therapeutic delivery across the brain parenchyma.

The disruption of the BBB contributes to both age-related vascular cognitive impairment^32^ and to neurodegenerative diseases^97^, including Alzheimer’s- and Parkinson’s disease^98^. This deterioration leads to elevated microvascular permeability, allowing harmful substances from the bloodstream to infiltrate the brain. These infiltrations contribute to neuroinflammation, oxidative stress, and neuronal damage^32^. We tested the beneficial effects of ApoE-FL/RSVs and FL/RSVs on age-related loss of BBB integrity. Treatments were conducted over both short (5 days) and long (1 month) durations. Surprisingly, both liposomal formulations significantly improved BBB integrity in the cortex of aged mice after only 5 days of treatment, which was evident in reduced microvascular permeability to both 40 kDa and 3 kDa molecular tracers. To compare liposomal delivery with oral RSV administration, we employed a longer treatment timeframe, as oral RSV has limited bioavailability and requires prolonged exposure to achieve therapeutic effectiveness. ApoE-FL/RSV delivery demonstrated remarkable efficiency. It produced near-complete restoration of BBB integrity in the aged mouse cortex after one month of treatment, as demonstrated by the full closure of the smallest molecular tracer in our *in vivo* permeability assay. This was accomplished with an almost 2000-fold lower RSV concentration than that achieved by oral delivery. The functional results were also supported by transcriptomic evidence for the beneficial effects of FL/RSV and ApoE-FL/RSV on BBB related gene sets. Molecular characterization of key BBB cell types, including endothelial cells, pericytes, and astrocytes, using scRNA-seq revealed that both ApoE-FL/RSV and FL-RSV treatments significantly enhanced cell–cell interaction strength and frequency in the aging brain. This suggests improved intercellular communication within the neurovascular unit. In addition, GO analysis demonstrated that both liposomal formulations effectively activated molecular pathways associated with BBB repair and maintenance, highlighting their potential to reverse age-related BBB dysfunction at the cellular and transcriptional levels.

The comparable short-term effects of FL/RSV and ApoE-FL/RSV on cortical BBB integrity can be explained by several factors. First, RSV may reach its maximal therapeutic effect at the concentrations delivered by both liposomal formulations, suggesting a dose-dependent saturation effect^99^. In this scenario, once the effective threshold for BBB rejuvenation is achieved, approaching levels observed in young animals, additional targeting by ApoE-FL does not confer added benefit, as the therapeutic ceiling has already been reached^99^. Second, although ApoE-FL and FL differ in their protein corona composition and targeting potential, they may have comparable RSV release profiles. If both formulations release RSV at similar rates, then differences in targeting efficiency would not significantly influence short-term therapeutic outcomes^100, 101^. Additionally, our assessment of BBB integrity was limited to cortical regions, which may not fully capture regional differences in therapeutic response. It is possible that deeper, subcortical structures, such as the hippocampus or thalamus, exhibit greater sensitivity to ApoE-FL/RSV due to their higher vulnerability to age-related vascular dysfunction and their distance to penetrating arterioles. Our measurements of vascular density in deep brain regions support this idea, showing more pronounced improvements with ApoE-targeted liposomal delivery. Nevertheless, future studies incorporating region-specific analyses will be critical to determine whether ApoE-mediated targeting confers superior benefits on BBB in anatomically distinct brain areas or under conditions of more severe cerebrovascular compromise.

Age-related cognitive decline is strongly linked to microvascular rarefaction, the progressive loss of microvessels in the brain^32^. This reduction in vascular density leads to impaired basal CBF, limiting the delivery of oxygen and nutrients to neurons, and contributing to neuronal dysfunction^32^. Compounding this is the uncoupling of NVC, the mechanism by which blood flow dynamically adjusts to meet the metabolic demands of active neurons. Impaired NVC exacerbates neuronal dysfunction, highlighting the critical importance of maintaining vascular integrity in aging brains to prevent or mitigate cognitive deficits^32^ ^9, 62, 102–105^. Restoring vascular density in aging brains, and consequently, basal CBF, holds significant clinical relevance. We found that both ApoE-FL-RSV and FL-RSV treatments fully rejuvenated vascular density in the aged cerebral cortex. However, in the hippocampus, only ApoE-FL-RSV treatment resulted in a significant increase in vascular density.

The reduced ability of FL-RSV to restore vascular density in the hippocampus compared with the cortex may reflect several factors. One key explanation is the regional heterogeneity of LRP1 expression across the brain^106,107, 108^. Studies have shown that cortical regions express higher levels of LRP1 than the hippocampus^84^, which likely facilitates more efficient receptor-mediated interactions even with FL-RSV nanoparticles that rely on their spontaneously formed, ApoE-enriched protein corona for endothelial targeting. In contrast, the hippocampus may have lower LRP1 expression, making ApoE-FLs, with their engineered ApoE protein corona, more efficient through stronger receptor engagement^109, 110^. Second, the hippocampus is also more vulnerable to vascular rarefaction during aging, with a 40% reduction in vascular density compared to 20% in the cortex, as our data show. This greater decline demands more precise and efficient targeting to achieve meaningful vascular restoration. The extent of age-related damage in the hippocampus may exceed the repair capacity of FL-delivered RSV. FLs, relying on a less stable and less consistent protein corona, may lack the precision and delivery efficiency needed to address this level of damage. Thus, achieving sufficient RSV concentrations in the hippocampus may require enhanced receptor-mediated docking and retention, provided by ApoE-FLs. These findings highlight the importance of targeted liposome engineering in addressing regional differences in vascular aging and ensuring effective therapeutic outcomes.

In addition to basal CBF, NVC plays a critical role in ensuring adequate neuronal function by dynamically matching blood supply to the metabolic demands of active neurons^111^. Neurovascular uncoupling is one of the early indicators of aging and is closely linked to cognitive deficits observed in aging^112^. Both ApoE-FL/RSV and FL/RSV treatments significantly improved cortical NVC in aging mice compared to aged-matched mice. However, the fact that only ApoE-FL/RSV achieved a level of cortical NVC comparable to that observed in young, healthy mice, suggests that the ApoE-functionalized liposomes were more effective, likely due to their enhanced targeting and retention in brain endothelial cells, which may lead to more precise or efficient delivery of RSV to the neurovascular units.

Resveratrol has shown some promising, but not always conclusive, effects on cognition^113^. While some studies suggest potential benefits in memory and cognitive function, particularly in animal models and specific populations, other studies, especially in humans, have yielded fewer positive results^114, 115^, often due to variations in dosing, timing, participant characteristics, and outcome measures. Additionally, the optimal dose and timing for cognitive benefits are still under investigation, making it difficult to draw firm conclusions^116,117^. Thus, the complete rejuvenation of spatial learning and memory function observed in our aged mouse model was striking, demonstrating a level of cognitive restoration rarely achieved in prior resveratrol studies^118, 119^ and underscoring the potential power of targeted cerebrovascular interventions. Importantly, as we did not find evidence for direct liposomal crossing into neurons, we conclude that the observed positive effects on cognition are primarily mediated through improvements in cerebrovascular function rather than through direct neuronal delivery of resveratrol.

In conclusion, this study introduces a biomimetic drug delivery paradigm that fundamentally shifts how cerebrovascular aging is targeted. By leveraging the naturally occurring ApoE-enriched protein corona of fusogenic liposomes, we demonstrate that full-length apolipoproteins can serve as self-assembling ligands for efficient, receptor-mediated brain endothelial delivery, without chemical modification or lysosomal trapping. This dual mechanism of ApoE-guided docking and membrane fusion enables unprecedented cerebrovascular specificity and intracellular drug availability, leading to complete rejuvenation of BBB integrity, vascular density, and neurovascular coupling in aged mice. These findings reposition the cerebrovascular endothelium as a primary therapeutic target for age-related cognitive decline and establish a versatile nanoplatform for vascular rejuvenation therapies.

While these findings provide strong evidence for the efficacy of ApoE-FL/RSV in rejuvenating the aged cerebrovasculature, several limitations should be noted. The study was conducted in aged mice, and translation to human physiology remains to be confirmed. In addition, detailed biodistribution and long-term safety studies are still required to assess potential off-target effects and optimize dosing for sustained therapeutic benefit.

## Materials and Methods

### Liposome composition

The phospholipids 1,2-dioleoyl-sn-glycero-3-phosphoethanolamine (DOPE) and 1,2-dioleoyl-3-trimethylammonium-propane (DOTAP) were purchased from Avanti Polar Lipids (Alabaster, AL, USA). The fluorescent dyes DiO, DiD, and DiR were purchased from Thermo Fisher Scientific (Waltham, MA, USA) and selected according to their spectral emission characteristics. RSV was obtained from Merck (Darmstadt, Germany) and Sigma Aldrich (St. Louis, MO, USA). Liposomes were hydrated in N-(2-hydroxyethylpiperazine-N-2-ethane sulfonic acid buffer (HEPES, pH 7.4, with 150 mM NaCl; Gibco, Waltham, MA, USA). For liposomal targeting human recombinant ApoE (PeproTech, Cranbury, NJ, USA, Cat. No. 350-02) was supplied in 10 mM sodium-bicarbonate buffer (1 mg/mL). All of the components were used without further purification. Unless otherwise stated, liposomes were formulated at a DOPE:DOTAP (10 mg/mL in chloroform) molar ratio of 1:1 (mol/mol), with a lipid-to-RSV (5 mg/mL in ethanol (EtOH)) ratio of 2:0.4, fluorescent dye (1 mg/mL in chloroform) incorporated at 0.05 molar equivalents relative to total lipid, and ApoE functionalization achieved at a lipid-to-protein molar ratio of 5000:1.

### APOE-FL/RSV/dye liposome preparation

All components were thoroughly mixed and dried under reduced pressure to form a lipid film. This dry film was then redispersed either directly in 20 mM HEPES buffer or in EtOH injected into 20 mM HEPES buffer to obtain FL at a final concentration of 10 mg/mL. The resulting particles were subjected to ultrasonic bath treatment (20 min at 4 °C) to homogenize their size. To generate ApoE-functionalized FL/RSV particles, liposomes were incubated with an ApoE stock solution at 20 °C for 1 hour with gentle agitation to allow spontaneous protein corona formation. FL/RSV liposomes can be stored at -20 °C for up to 6 months, while FL/RSV and ApoE-FL/RSV can be stored at 4 °C for up to 2 weeks. Stocks can be directly diluted in 1× phosphate buffered saline (PBS) to the desired concentration. For *in vivo* and *ex vivo* experiments liposomes were used at 10 mg/mL concentration, while *in vitro* and *in vivo* biological effectiveness experiments used 2 mg/mL concentration. Detailed description of concentration, injection volumes, frequencies, and treatment durations are provided in the respective experimental section.

### Analysis of protein corona composition on FL/RSV particles using LC-MS/MS

FL/RSV (2 mg/mL) particles were incubated with human plasma (HP) or mouse plasma samples (Merck) for 1 h at 37 °C with gentle agitation. Protein aggregates were isolated by 30 min centrifugation (16.1 krcf, 4 °C). After removal of the supernatant the HP-FL/RSV pellet was washed with PBS and redispersed in 1.5% sodium dodecyl sulphate (SDS) solution. Proteins were denatured for 1 h at 95 °C and stored at -20 °C until further use. For LC-MS/MS sample preparation 10 µg of protein were reduced with dithiothreitol (10 mM; 15 min, 95 °C) and alkylated with chloroacetamide (25 mM; 30 min). Alkylation was stopped by adding 50 mM dithiothreitol (20 min). For buffer exchange, single-pot solid phase (SP3) paramagnetic beads were used^120^. Proteins were bound to the beads by addition of four sample volumes EtOH (80% final concentration), washed twice with 90% acetonitrile (ACN), and released by reconstitution in digestion buffer containing 50 mM HEPES buffer (pH 7.5, 2.5 mM CaCl_2_). Trypsin was added at a protein:protease ratio of 100:1 (w/w) and incubated for 18 h at 37 °C. The samples were acidified to pH<3.0 by addition of 1% formic acid (FA). Peptide purification was achieved using a stage tip approach following the example of Rappsilber *et al.* using double packed cation exchange layers (Merck)^121^. Peptides were eluted with 5% ammonia solution (NH_4_OH) in 60% ACN.

Desalted peptide solutions were concentrated under reduced pressure, taken up in 2% ACN with 0.1% FA, and injected into an UltiMate 3000 RSLCnano system (Thermo Fisher Scientific), equipped with µPAC trap and analytical (50 cm flow path) reversed phase columns (PharmaFluidics, Ghent, Belgium). Separated peptides were ionized using a CaptiveSpray nanoBooster ion source (Bruker, Billerica, MA, USA) and introduced into an Impact II quadrupole time-of-flight tandem mass spectrometer (Bruker) as described^122^. Using version 5.1 of Brukeŕs HyStar software, MS data were acquired in line-mode working at a mass range from 200 to 1750 m/z and an acquisition rate of 5 Hz. The 14 most intense ions were selected for fragmentation, with fragment spectra automatically being recorded at 5 to 20 Hz depending on the precursor intensity. Selected precursors were excluded for the next 0.4 min unless signal to noise ratio improved 3-fold.

Data were processed in MaxQuant software (v2.0.1.0, Max Planck Institute of Biochemistry, Martinsried, Germany)^123^ using the UniProt mouse or human database^124^. Protein quantification within a sample was based on the intensity-based absolute quantification (iBAQ) algorithm. The algorithm provides a quantity mapping iBAQ value for each protein. The entirety of all iBAQ values formed the basis for further analysis and was used to determine a sample’s protein composition. According to the formula below, the molar fraction of a specific protein was obtained by dividing the protein’s iBAQ value by the sum of all iBAQ values. The resulting relative iBAQ value (riBAQ) is a normalized measure for a protein’s molar abundance within the analyzed protein mixture.

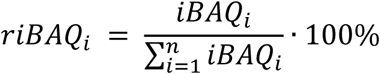

### Particle characterization via dynamic and electrophoretic light scattering (DLS, ELS)

Hydrodynamic diameter and PDI were determined by DLS using a Zetasizer Nano ZS (Malvern Panalytical, Malvern, UK) at a scattering angle of 173° and a temperature of 22 °C. Briefly, FL stocks (2 mg/mL) were diluted 1:10 in 20 mM HEPES buffer (pH 7.4). 50 µL of diluted samples (n=11) were pipetted into a SUPRASIL quartz precision cuvette with a path length of 3 mm (Hellma Analytics, Müllheim, Germany). Zeta potential was also measured using ELS applying the Smoluchowski approximation. Samples from the hydrodynamic diameter measurements were diluted 1:15 in 20 mM HEPES (pH 7.4). 750 µL diluted samples (n>3) were transferred to a DTS1070 disposable capillary cell (Malvern Panalytical) and measured at 22 °C using 20 individual scans. Data evaluation was done using version 7.13 of Malvern’s Zetasizer software.

### ApoE-FL stability study in plasma

Plasma was obtained from fresh human or mice blood collected in lithium heparin tubes (Sarstedt, Nümbrecht, Germany) and mixed gently to avoid clotting. Afterwards, blood was centrifuged (10 min, 2 krcf, RT) and the plasma was carefully removed with a pipette and centrifuged again (30 min, 16.1 krcf, 4°C). The isolated human or mice plasma was mixed with ApoE-FL/RSV stocks (2 mg/mL) at a ratio of 4:1 (v/v). Multiple formulations featuring different lipid/ApoE ratios (10000/1, 5000/1 and 1000/1 (mol/mol)) were tested. In addition, an ApoE-only reference control was prepared by mixing ApoE in HEPES with rat plasma at a ratio of 4:1 (v/v). Samples were incubated at RT with gentle shaking and aliquots were collected after ¼, 1, 3 and 24 h and centrifuged immediately (30 min, 16.1 krcf, 4 °C). The ApoE content in the lipid-free supernatant was quantified using a human ApoE ELISA kit (Thermo Fisher Scientific). Before the assay, samples were diluted according to the expected ApoE quantity in order to remain within the detection range of the kit. The kit was used as recommended by the manufacturer and the absorbance was recorded at 450 nm using an Infinite M1000 PRO plate reader (Tecan, Männedorf, Switzerland). The FL-bound ApoE proportion was calculated by comparison of the ApoE amounts identified in the supernatants and the corresponding lipid-free reference samples.

### Morphometric analysis of the liposomes using Cryo-TEM

The 200 mesh R2/1 holey carbon Quantifoil grids (Electron Microscopy Sciences, Hatfield, PA, USA) were glow discharged in a Pelco easiGlow Glow Discharge Cleaning System (Ted Pella Inc, Redding, CA, USA) for 90 s at 15 mA and a 15 s hold time. Freshly prepared liposome suspensions were vitrified in liquid ethane using an EM GP2 Automatic Plunge Freezer (Leica Microsystems, Wetzlar, Germany) at 10 °C chamber temperature and 80% relative humidity. 4 µL sample volume was applied to the front of the grid and blotted from the backside for 3–4 seconds. Next, samples were transferred under cryo-conditions to a 200 kV Talos Arctica G2 transmission electron microscope (Thermo Fisher Scientific), equipped with a Bioquantum GIF (Gatan, Pleasanton, CA, USA) and a K3 direct electron detector (Gatan). Approximately 200–400 micrographs were recorded per sample at a magnification of 31,000×, using a 20eV energy filter slit and with the K3 camera in super-resolution mode. Micrographs were recorded as movies with 12 frames with a dose of ∼1 e^-^/Å^2^/frame for a cumulative dose of 12.7 e^-^/Å^2^/micrograph, a nominal defocus of -6 µm and a final pixel size of 0.28 Å/px. All micrographs were motion and contrast transfer function corrected in WARP^125^. Micrographs showing clear sample features were analyzed using CryoVIA^126^. After the membranes were detected by the default pre-trained neural network, the dataset was manually curated, and obvious liposomes were removed from the detected nanodiscs. Diameters were measured as the greatest distance between two points on segmented contour, measured at the point in the center between the two phospholipid leaflets. Measurements were exported from CryoVIA and analyzed in OriginPro 2023 (version 10.0.0.154; OriginLab Corporation, Northampton, MA, USA)

### Assessment of RSV encapsulation and ApoE binding efficiency

FL/RSVs with varying lipid/ApoE ratios containing DOPE, DOTAP, DiO, and RSV at a molar ratio of 1/1/0.05/0.4 (mol/mol) were prepared as described previously. Following 1 h incubation with ApoE, 25 µL of each liposome stock (2 mg/mL) were diluted 1:4 (v/v) with 20 mM HEPES buffer (pH 7.4). Diluted samples were transferred into Nanosep® centricon tubes (molecular weight cut-off 100 kDa) (Pall Corporation, Port Washington, NY, USA) and centrifuged for 30 min (5 krcf, 4 °C) to separate unencapsulated RSV. The filtrate was collected, diluted 1:4 (v/v) with PBS, and loaded into a SUPRASIL® quartz glass cuvette with a 10 mm path length (Hellma Analytics). Quantification based on intrinsic fluorescence of RSV upon excitation at 330 nm using a Fluorolog-3 spectrometer (HORIBA Jobin Yvon, Oberursel, Germany) operated with FluorEssence (v3.8.0.60) software. Emission spectra were collected from 350–600 nm and baseline corrected by subtracting the solvent spectrum. The RSV concentration was calculated based on a calibration curve determined in PBS buffer using the spectrás maximum intensity at 395 nm. The encapsulation efficiency was assessed by dividing the amount of FL bound RSV by the corresponding RSV input.

To assess ApoE binding, the same FL/RSV preparation used for the RSV encapsulation assay was processed, and the resulting filtrate was diluted 1:4 (v/v) and tested for unbound ApoE using a human ApoE ELISA kit (Thermo Fisher Scientific). The assay was performed according to the manufactureŕs manual. Absorbance measurements at 450 nm were conducted on an Infinite M1000 Pro plate reader (Tecan) using i-control software (v3.9.1.0). Bound ApoE was calculated by subtracting the concentration of ApoE detected in the filtrate from the total ApoE input for each lipid/ApoE ratio.

### Cell cultures and generation of LRP1 KD in vitro model using CRISPR/Cas9

Human cerebral microvascular endothelial cell line (hCMEC/D3, Sigma Aldrich) was used as an *in vitro* endothelial model for all mechanistic and trafficking/targeting assays. Cells were grown in a flask pretreated with an attachment factor solution (Cell Systems, Kirkland, WA, USA) for 30 minutes at 37 °C. Then, cells were expanded in Complete Classic Medium with serum (Cell Systems) supplemented with CultureBoost™, Penicillin-Streptomycin (5 mL, 10,000 U/mL, Gibco), and Bac-Off Antibiotic (1mL, Cell Systems). Cultures were maintained under standard endothelial cell conditions (37 °C and 5 % (v/v) CO_2_ in a humidified atmosphere).

To determine whether LRP1 plays a pivotal role in ApoE-FL uptake and affects fusion capacity, a stable LRP1 KD hCMEMC/D3 line was generated using CRISPR Cas9/GFP-based genome-editing system containing GFP reporter. Cells were seeded at a density of 100,000 cells per well in 6-well plates (Thermo Fisher Scientific) and grown to 60% confluence. After 24 hours, LRP1-targeting CRISPR sgRNA AAV (saCas9; Serotype 9; Applied Biological Materials Inc., Richmond, BC, Canada) was mixed with Complete Classic Media (200 μL virus/6 mL) and 1 mL of this master mix was applied for each well. 72 hours after viral delivery, cultures were harvested, GFP-positive cells were enriched using fluorescence-based cell sorting (Wolf Cell Sorter, Nanocellect, San Diego, CA, USA) and subsequently re-expanded. This enrichment process was repeated to obtain a stable population with consistent reporter expression. Flow-cytometric analysis (Guava EasyCyte BGR HT, Cytek Biosciences, Fremont, CA, USA) was used to monitor GFP enrichment. Both the gene-edited LRP1 KD and wild type hCMEC/D3 lines were subsequently used for comparative experiments assessing the LRP1 dependence of ApoE-FL uptake and fusion behavior.

### Immunofluorescent detection of ApoE on hCMEC/D3 cells following liposomal treatment

HCMEC/D3 cells were seeded on 8-well µ-slides (ibidi, Graefelfing, Germany) for 72 h under standard conditions discussed previously. Prior to liposomal treatment, FL/RSV or ApoE-FL/RSV at a lipid/ApoE ratio of 5000/1 and 1000/1 mol/mol and a lipid concentration of 2 mg/mL were prepared freshly and diluted 1:20 (v/v) with cold PBS. Culture medium was aspirated, and 200 µL of liposome stock solution was added to each chamber. After 20 min incubation at 37 °C, the liposome solution was replaced by 250 µL of fresh Endothelial Cell Growth Medium MV2. As for control, cells were incubated with recombinant ApoE solution (2 µg/mL in PBS) equivalent to the highest ApoE concentration in FL samples. Afterwards, cells were fixed with 3.7% (w/w) paraformaldehyde (PFA) in PBS for 15 min at RT followed by three PBS washes. Blocking was performed with 5% (w/w) milk powder in PBS for 1 h at RT. After another washing procedure, the antibody stock was prepared by diluting CoraLite Plus 488-conjugated anti-ApoE antibody (Thermo Fisher Scientific) 1:100 (v/v) with 1% (w/w) milk powder in PBS, then antibody stock was added to each chamber and cells were incubated for 2 h at RT. Stained cells were washed three times with PBS for 5 min on an orbital shaker. Nuclei were counterstained by adding one drop of NucBlue reagent for fixed cells (Thermo Fisher Scientific). Fluorescent imaging was performed on a LSM710 confocal laser scanning microscope (Carl Zeiss Microscopy, Jena, Germany) equipped with an EC Plan-Neofluar 40x/1.30 Ph3 oil-immersion objective (Carl Zeiss Microscopy) and an XL-2 stage incubator (37 °C and 5% CO_2_ (v/v)). The system was operated using ZEN black software (v14.0.7.201, Carl Zeiss Microscopy). CoraLite Plus 488-conjugated ApoE antibody was excited using an argon ion laser (488 nm) and detected with a band pass filter BP 500–555 nm. DiR excitation was achieved using a helium-neon laser (633 nm) and the emitted light was collected with a long pass filter LP 660 nm. NucBlue was excited by a 405nm laser diode and detected with BP 415–475 nm filter. Micrographs were recorded with a resolution of 13 pixel/µm. Image processing was performed in Fiji^127^ (ImageJ platform v2.14)^128^, where brightness and contrast of individual channels were adjusted to ensure optimal visualization.

### In vitro colocalization and quantitative analysis of liposomal uptake and subcellular trafficking

hCMEC/D3 cells were seeded on 8 well µ-slides (ibidi) and cultivated for 72 h until forming a confluent monolayer. To visualize the subcellular localization of FL, cells were labeled with organelle-specific fluorescent markers. ER-Tracker Green, LysoTracker Green DND-26, MitoTracker Green FM, and CellMask Green were used to label endoplasmic reticulum, lysosomes, mitochondria or plasma membrane, respectively. All reagents were purchased from Thermo Fisher Scientific and applied as 1 mM stock solution in dimethyl sulfoxide. Each stock solution was diluted 1:1000 (v/v) in culture medium before single organelles were stained by removing the medium and incubating the cells with the respective staining solution (1 hour, 37 °C). After staining, hCMEC cells were incubated with FL (DiR) solution and imaged as previously described.

Confocal microscopy images from the previously described *in vitro* experiments were acquired as z-stacks containing multiple fluorescence channels. The image stacks were separated into individual channels for further analysis. Image stacks from the GFP-positive LRP1 KD CRISPR/Cas9 cells were converted to 8-bit grayscale, and then the Trainable Weka Segmentation plugin^129^ was employed on the green channel to segment GFP-positive cell profiles only. Following segmentation, Otsu thresholding was applied to generate a binary mask which was inverted to exclude GFP-negative cellular regions from the analysis. For the wild type hCMEC/D3 cells lacking GFP-expression, the green channel was removed entirely and the whole field of view was used. The remaining channels were converted to 8-bit grayscale and subjected to a rolling ball background subtraction (radius of 5 pixels) across the z-planes. The background-corrected channels were binarized using the Otsu method. To assess the colocalization between the liposomal dye and the different organellar markers, the built-in image calculator function of Fiji was used to perform a logical ’AND’ operation between the relevant channels to identify overlapping regions where both markers were present. The resulting overlap images were analyzed z-plane by z-plane to calculate the percentage of pixels with non-zero intensity across the z-stack, representing the degree of colocalization without introducing a projection bias by merging z-planes into a two-dimensional projected image. A similar measurement was performed on the liposome channel alone to quantify its distribution.

### Analysis of mitochondrial oxidative stress reduction following ApoE-FL/RSV treatment

hCMEC/D3 cells were cultivated as previously described and seeded on 8-well µ-slides (ibidi) for mitochondrial stress assays. Mitochondrial stress was induced by incubation in high concentrated glucose solutions (50 and 150 mM) for 48 h. Subsequently, cells were treated with ApoE-FL/RSV solution (lipid/ApoE 5000/1 (mol/mol) and lipid/RSV 2/0.4 (mol/mol)) at final concentration of 0.1 mg/mL for 20 min. After treatment, liposome solution was replaced by fresh cell culture medium, and cells were maintained for an additional 24 h under standard cell culture conditions. Cells were subsequently detached using trypsin/EDTA solution (Thermo Fisher Scientific), resuspended in 300 µL culture medium containing 2% (v/v) fetal calf serum (Thermo Fisher Scientific) and incubated with a mitochondrial membrane potential dye, JC-1 (Thermo Fisher Scientific), at a concentration of 2 µM for 25 min at 37 °C. Afterwards, cells were centrifuged (600×g, 3 mins) and resuspended in PBS. Fluorescence intensities of JC-1 aggregates (red, excitation at 488 nm, emission at 585 nm) and monomers (green, excitation at 488 nm, emission at 525 nm) were measured on a CytoFLEX S flow cytometer (BC Life Sciences, Indianapolis, IN, USA). Untreated cells served as baseline controls. Flow cytometric data were analyzed using CytExpert software (BC Life Sciences).

### Quantification of LRP1 expression at protein and transcript level using western blot and RT-qPCR

LRP1 expression was evaluated *in vitro* using lysates from GFP-positive LRP1 KD and wild type hCMEC/D3 cells, and *in vivo* using cortical tissue samples from mouse brains described below. Cells (5×10^6^) and tissue samples (40 mg) were homogenized in 1 mL of RIPA lysis buffer (Thermo Fisher Scientific), then centrifuged at 16 krcf for 20 minutes. To increase the protein yield, the pellet was sonicated for 30 seconds at 50% pulse intensity with a Cl-18 probe sonicator (Thermo Fisher Scientific), in an ice bath. Total protein concentration from the supernatant was quantified using the bicinchoninic acid (BCA) protein assay (Thermo Fisher Scientific). LRP1 protein abundance was quantified using Anti-Rabbit Detection Module (Biotechne, Minneapolis, MS, USA) against rabbit anti-LRP1 monoclonal antibody (Abcam, Cambridge, UK) with JESS Automated Western Blot System (Biotechne, Minneapolis, MN, USA) based on the manufacturer’s recommendation. For signal processing and analysis, the pre-established pipeline, set by the manufacturer was used in the JESS Compass software (v6.1.0, Biotechne).

LRP1 transcript level was assessed from brain tissue sampled using RT-qPCR. Samples were collected and homogenized in Buffer RLT lysis buffer (Qiagen, Germantown, MD, USA). Then total RNA was extracted from tissues using RNeasy Mini RNA purification kit (Qiagen) in QIAcube Connect automated system (Qiagen). RNA concentration and purity were assessed using NanoDrop UV-Vis Spectrophotometer (Thermo Fisher Scientific). cDNA was synthesized from 1 µg of total RNA with the High-Capacity RNA-to-cDNA Kit (Thermo Fisher Scientific), according to the manufacturer’s instructions. RT-qPCR was carried out on an Applied Biosystems QuantStudio 12K Flex system (Thermo Fisher Scientific) using a TaqMan gene expression assay for LRP1 (Thermo Fisher Scientific). Cycling conditions consisted of an initial hold at 50 °C, denaturation at 95 °C, and annealing/extension at 60 °C for 40 amplification cycles. Relative gene expression levels were calculated using the 2^ΔΔCt^ method, with β-actin serving as the endogenous control for normalization.

### Animal models and experimental conditions

C57/BL6 and Rosa26-tdTomato transgenic mice of different age groups were used to investigate the biodistribution and functional impact of RSV-loaded FL. For quantification of *in vivo* liposomal deposition 3–9-months old young adult mice were selected to minimize age-related autofluorescence. Functional studies including, the assessment of NVC, microvascular density, BBB integrity, and other endothelial physiological assessments and cognitive performance have been executed on animals from different age-groups from young adults (3–9-month-old) to aged (> 18-month-old).

Animals were housed under standardized environmental conditions with 12-hour light-dark cycle, and *ad libitum* access to water and a standard AIN-93G diet. Initially, the mice were housed in a specific pathogen-free environment within the Rodent Barrier Facility at the University of Oklahoma Health Sciences Center (OUHSC) and subsequently transferred to the standard rodent colony at OUHSC for the experimental procedures. All experiments were conducted in strict adherence with the ethical guidelines of the National Institutes of Health (NIH) Guide for the Care and Use of Laboratory Animals (NIH Publications No. 8023, revised 1978) and were approved by the Institutional Animal Care and Use Committee at OUHSC. Throughout the study, measures were taken to minimize potential discomfort and ensure optimal animal welfare.

### In vivo treatment paradigms for retro-orbital liposomal and oral RSV delivery

For *in vivo* administration of FL/RSV and ApoE-FL/RSV formulations, mice received liposomes via retro-orbital injection under inhalational anesthesia with 3% isoflurane for induction and 2% during the procedure. Liposomes were prepared and stored as described previously and the injection volume and dosage varied by experimental purpose and adjusted according to the liposomal concentration. High concentration liposome preparations (10 mg/mL) were used for studies focusing on liposome visualization, including tracking and biodistribution. For assessing biological effects (NVC, BBB, RAWM, and *ex vivo* aorta ring experiments), a lower concentration (2 mg/mL) was applied. The specific dye concentration of liposomes, volume, frequency, and duration of injections are detailed within each respective study section. Oral RSV was administered in experiments assessing the biological effect on NVC, BBB permeability, RAWM performance, and *ex vivo* aorta function to serve as a classical route of administration. RSV was dissolved in 100% EtOH at a concentration of 60 mg/mL and incorporated into a strawberry-flavored, sugar-free gelatin vehicle (Jell-O, Walmart). The gelatin mixture was poured into molds containing 24 mg RSV, then they were left to set at 4 °C, overnight. The gelatin RSV molds were placed in cages of 4 mice; therefore, each mouse received approximately 6 mg RSV/day. Gelatin molds were administered every weekday for the duration of the treatment period, ranging from 1 to 3 months depending on the experimental setup.

### Quantification of transient interaction of fusogenic liposomes with circulating and splenic immune cells

Liposome uptake by circulating immune cells was assessed following retro-orbital injection of FL or ApoE-FL liposomes (30 µL, 2 mg/mL). Before injection and at 10-, 30-, and 60-minute post-injection, whole blood was collected via submandibular venipuncture in the heparin-coated tubes, and 10 µL of blood was diluted in 990 µL of MACS buffer (Miltenyi Biotec, Bergisch Gladbach, Germany). The suspension was passed through a 40 µm mini cell strainer (PluriSelect, Leipzig, Germany) and centrifuged at 500 rcf for 10 min. The supernatant was discarded, and the cell pellet was resuspended in 1 mL MACS buffer, followed by a second centrifugation at 500 rcf for 10 min. The final pellet was resuspended in 200 µL MACS buffer, and the cell suspension was transferred to a 96-well plate for flow cytometry analysis.

Liposome uptake by the spleen was assessed following retro-orbital injection of FL or ApoE-FL liposomes (30 µL, 2 mg/mL). At 2 h, 6 h, and 24 h post-injection, mice were euthanized and perfused with ice-cold PBS. After 12 minutes of perfusion, spleens were collected and finely minced on a petri dish, transferred to 50 mL tubes, and centrifuged at 50 rcf for 5 min at 4 °C. Samples were washed with 5 mL of sterile PBS, after which digestive enzymes (50 µL collagenase, 2 µL elastase, 2 µL dispase, and 3 µL hyaluronidase) were added. Tissues were incubated in a rotating incubator at 37 °C for 10 min. Following enzymatic digestion, samples were gently dissociated by pipetting first with a 10 mL serological pipette, then with a 5 mL pipette. The resulting cell suspension was filtered sequentially through 100 µm and 30 µm strainers. Cells were centrifuged (300 rcf, 10 min, 4 °C) and resuspended in 1 mL PBS. Samples were further centrifuged at 1000 rcf for 10 min at 4 °C. The supernatant was discarded, and the pellet was resuspended in 1 mL of MACS Buffer for flow cytometry. Data was acquired using a Guava EasyCyte BGR HT (Cytek Biosciences) flow cytometer. For each sample, 10,000 peripheral blood cells and 20,000 spleen cells were analyzed in duplicates. Liposome interaction was assessed based on the DiO positivity of analyzed cells. The DiO+ gate was defined using baseline green fluorescence for blood and spleen cells determined separately via a fluorescence-minus-one approach with pre-injection samples. Liposome uptake was expressed as the percentage of DiO+ cells.

### Sample preparation for scRNA-seq and transcriptomic data analysis

Mice were anesthetized with isoflurane and transcardially perfused with ice-cold PBS. Brains from young (N=5) and aged mice (N=5) served as controls, and additional aged cohorts received FL/RSV, ApoE-FL/RSV, or orally administered RSV (N=3 per group). Following perfusion, whole brains were rapidly collected and dissected into small pieces. After a brief spin down (5 min, 50 rcf) tissue fragments were subjected to enzymatic dissociation in PBS using a defined mixture of collagenase, hyaluronidase, elastase, and dispase (0.1%, 0.06%, 0.04% and 0.04% (v/v)) under controlled agitation conditions (45 min, 37 °C). The resulting cell suspensions were passed through 70-µm and 30-µm mesh filters (Miltenyi Biotec) and subsequently processed with Debris Removal Solution (Miltenyi Biotec) to reduce myelin and other contaminants. Cell pellets were resuspended in PBS containing 0.1% BSA and stained with SYTOX Green Nucleic Acid Stain (Invitrogen, Waltham, MA, USA) to identify viable cells for purification on a Wolf Cell Sorter (NanoCellect). Sorted live cells were collected, washed, and resuspended in a minimal volume of 0.04% BSA in PBS solution according to yield. High-quality live cells were used to prepare single-cell cDNA libraries with the Chromium Single Cell 3′ platform (10x Genomics, Pleasanton, CA, USA), following the manufacturer’s guidelines. Libraries were generated using the Chromium Single Cell 3′ Chip and Library & Gel Bead Kit v2 and sequenced on an Illumina NovaSeq 6000 system (San Diego, CA, USA).

Raw fastq files were quality checked and processed using Cell Ranger (10x Genomics) to generate aligned and demultiplex count files. Filtered gene-barcode matrices were loaded into Seurat, and data were normalized with a scale factor of 10,000. Highly variable genes were identified and used for scaling and principal component analysis. A shared nearest-neighbor graph was constructed, and clusters were identified using Seurat’s graph-based clustering, followed by Uniform Manifold Approximation and Projection (UMAP) were used for visualization. Putative doublets were detected with DoubletFinder (expected rate 8.7%) and excluded from downstream analyses. Cell-type labels were assigned using scMRMA and further refined by manual inspection of canonical marker genes. Endothelial cells were bioinformatically subsetted and reclustered. For each annotated cell type, marker genes were ranked by log fold-change, and the top 20 were used to define gene signatures as previously described^130^. AUCell was used to calculate per-cell area under curve (AUC) scores for each signature, and cells were assigned to the signature with the highest AUC score, provided the maximum exceeded 0.25; cells below this threshold were manually curated using known markers. Interactive visualization was performed using ShinyCell, and intercellular communication networks were inferred with CellChat based on ligand–receptor interactions between annotated cell types.

### Assessment of cognitive function using the RAWM

To assess the impact of RSV delivery system/approach on cognitive function, learning, spatial memory and long-term memory, comparing traditional oral administration with advanced liposomal delivery systems. C57BL6 mice were assigned as young controls (5 months, N=15), aged controls (19 months, N=15); FL/RSV-treated (18 months at treatment onset, N=15) and ApoE-FL/RSV-treated (18 months at treatment onset, N=15) aged groups. Liposomal treatment was used via retro-orbital injections, and oral RSV was administered in gelatin vehicle. Liposomal formulations containing DiO were administered according to the dosing paradigms described previously. Oral RSV was provided daily on weekdays over a one-month period.

After the treatment, cognitive function, spatial memory and long-term memory were assessed using the RAWM, following our published protocols^65^. Data were analyzed using Ethovision software (Noldus Information Technology, Leesburg, VA, USA). Each daily session involved two blocks of four consecutive acquisition trials, during which the mice learned the location of the submerged escape platform. Starting from an arm without the platform, the mice had up to one minute to find it and were allowed to remain on the platform for 15 seconds after each trial. The platform’s location remained consistent throughout trials. Over the course of three days, the mice showed gradual improvements in performance. Errors were recorded when a mouse entered the wrong arm, defined by having all four paws in the distal half of the arm. Once the group acquired the task, the mice were returned to their home cages for seven days, followed by a recall/probe trial on day 10. On day 11 (reversal/extinction phase), the mice were tested on their ability to relearn the task with the platform moved to a new arm, neither adjacent to nor directly opposite the original location.

### Assessment of endothelium-dependent vasorelaxation in aortic ring preparations

These experiments aimed to evaluate the effects of RSV on systemic endothelial function, comparing traditional oral administration with advanced liposomal delivery systems using the aorta ring vasorelaxation assay on C57BL6 mice were assigned as young control (6 months, N=3); aged control (24 months, N=3); FL/RSV-treated (20 months at treatment onset, N=3) or ApoE-FL/RSV-treated (20 months at treatment onset, N=3) aged groups. Following the previously described treatment paradigms, endothelium-dependent vasorelaxation was evaluated using isolated aortic ring preparations as previously described^27^. Briefly, aortas were sectioned into 1.5 mm ring segments and mounted in myograph chambers (Danish Myo Technology, Hinnerup, Denmark) to measure isometric tension. The vessels were perfused with Krebs buffer solution (118 mM NaCl, 4.7 mM KCl, 1.5 mM CaCl_2_, 25 mM NaHCO_3_, 1.1 mM MgSO_4_, 1.2 mM KH_2_PO_4_, and 5.6 mM glucose at 37 °C, gassed with 95% air and 5% CO_2_). After a 1 h equilibration period optimal passive tension was established, then the aortic rings were pre-contracted with 10^−6^ M phenylephrine hydrochloride (Sigma Aldrich) in Krebs buffer solution. Endothelium-dependent relaxation was assessed by applying acetylcholine chloride (Sigma Aldrich) across a graded series of concentration range from 10^-4^ M to 10^-9^ M diluted in Milli-Q water.

### Real-time monitoring of FL/RSV and APOE/FL/RSV with intravital two-photon microscopy

#### Chronic cranial window surgeries

Chronic cranial windows were prepared as described in previous publications^9, 15, 62, 65, 74, 131^. Briefly, mice were anesthetized with 2–3% isoflurane and secured in a stereotactic stage under a Stemi 2000 stereomicroscope (Carl Zeiss Microscopy). After preparation of the surgical field, craniotomy was performed over the somatosensory cortex. Then the bone fragment was removed, and a glass coverslip (Warner Instruments, MA, USA) was sterilized and placed over the exposed cortical surface. Cyanoacrylate glue was applied around the edges of the coverslip to secure it to the bone, followed by a layer of acrylic cement (Stoelting, Wood Dale, IL, USA) applied around the window and on the exposed skull. Afterwards, buprenorphine (1 mg/kg, sc) for pain relief and enrofloxacin (10 mg/kg, sc) to prevent infection were administered and mice were monitored during the recovery. All imaging experiments were performed at least 2 weeks post-surgery.

#### In vivo two-photon imaging of liposomes

C57BL6 mice with chronically implanted cranial window were treated with FL/RSV (5 months, N=5) or ApoE-FL/RSV (5 months, N=10) and used to assess the real-time adhesion of the previously described liposomes to cerebral vessel walls using intravital two-photon microscopy.

For imaging, mice were anesthetized and positioned in a stereotaxic frame under a DIVE multiphoton microscope (Leica Microsystems) equipped with InSight X3 tunable two-photon laser (Spectra-Physics, Milpitas, CA, USA). Fluorescent signal from the liposomes was visualized using 800-nm excitation and the emitted light was detected by non-descanned detectors with three filter settings (480–550, 630–740, and 760–795 nm). Cerebral vasculature was labeled with 500 kDa FITC-dextran tracer (4 mg/mL, Sigma Aldrich). After baseline acquisition, 120 µL of 10 mg/mL of DiD-labeled FL/RSV or ApoE-FL/RSV was injected retroorbitally. Imaging was performed at five different cortical locations across multiple time-points (pre-injection, 5 min, 30 min, 1 h, 24 h, 48 h post-injection) with a 25× HC FLUOTAR L 25x/0,95 W VISIR water immersion objective (Leica Microsystems).

Recorded z-stacks were exported, and analyzed in the IMARIS (v10.2, Oxford Instruments, Abingdon, UK). Vascular volumes were reconstructed from the FITC-dextran channel using the built-in multiscale 3D intensity-based segmentation framework, which applies hierarchical smoothing and voxel-level classification to separate vessels from background signals. Liposomal signals were identified using the built-in spot/object detection module with morphology- and intensity-based thresholds optimized for small particles. Following detection, 3D object segmentation was refined using surface-rendering algorithms that incorporate local contrast, edge gradients, and size constraints to resolve individual liposome structures within dense vascular regions. To categorize intravascular particle behavior, segmented liposomes were classified using the built-in supervised machine learning workflow in IMARIS, which incorporates object-level features such as displacement vectors, shape descriptors, elongation, and orientation. This enabled robust discrimination between stationary, vessel-adherent liposomes and freely circulating liposomes.

### BBB permeability assessment with intravital two-photon microscopy

The effect of RSV on BBB permeability in aging mice was assessed by comparing oral administration with intravenous FL delivery systems. C57BL6 mice with chronic cranial window were used as young controls (5 months, N=9), aged controls (24–26 months, N=9), FL/RSV-treated (25 months, N=8), and ApoE-FL/RSV-treated (25 months, N=8) aged groups. Treatment groups received liposomes for four consecutive days, and BBB permeability was measured on the fifth day. To assess the long-term effect of RSV another five groups of young control (5 months, N=9), aged control (19–22 months, N=20), oral RSV-treated (18 months, N=5), FL/RSV-treated (18 months, N=5), and ApoE-FL/RSV-treated (18 months, N=5) aged groups were treated with orally administered RSV every weekday for one month. Treatment with the previously described liposomal formulations was administered via retro-orbital delivery for five consecutive days, followed by twice-weekly injections for the remainder of the one-month treatment period. Mice were anaesthetized and prepared for imaging as previously described. Afterward, the stereotaxic stage was transferred under the Fluoview FV1000 two-photon microscope (Olympus, Tokyo, Japan), equipped with a 25× XLPLN25XWMP water immersion objective (Olympus) and an 800-nm laser. Emitted light was detected by PMT detectors using three filter sets (420–460, 495–540, and 575–630 nm). 500kDa FITC-dextran tracer (2mg/mL in sterile saline, Thermo Fisher Scientific) was retro-orbitally injected to label the vasculature and establish a baseline. Subsequently, fluorescent tracers with decreasing molecular weight were injected starting with a 40 kDa FITC-dextran, followed by a 3 kDa FITC-dextran (both from Thermo Fisher Scientific). Finally, sodium fluorescein (Sigma-Aldrich) was administered as a 0.3 kDa tracer. After each injection, a time-Z-stack over 15 minutes was acquired, resulting in a hyperstack.

BBB permeability was quantified using a “relative permeability changes over-time” paradigm. For each tracer concatenated time-Z-stacks were processed by subtracting the baseline maximal intensity z-projection (I_0_ [a.u.]) from subsequent frames to isolate extravascular fluorescence increase. Vascular masks were generated using the “Trainable Weka Segmentation” plugin of Fiji and refined by subtracting local autofluorescence to accurately extract the parenchymal signal. Changes in tracer signal intensity (I [a.u.]) were expressed relative to the baseline (I_0_). The cumulative intensity-time profiles were integrated to obtain the area under the curve that served as the metric of relative BBB permeability within the imaged volume.

### Assessment of liposomal biodistribution in different tissues

To assess the biodistribution of FL/RSV and ApoE-FL/RSV in 5-month-old male C57BL6 mice were used. Experimental groups included untreated controls (N=5), FL/RSV-treated (N=7), and ApoE-FL/RSV-treated (N=7) mice. The treatment groups received 120 µL of 10 mg/mL DiD-labeled liposome solution, while control mice received no injection. Animals were anesthetized with isoflurane, transcardially perfused after 30 minutes post-injection with 1× PBS at 110 mmHg for 10 minutes, followed by an additional 50 mL of 4% PFA. Organs analyzed for biodistribution included the brain, liver, kidney, and quadriceps. After postfixation for 24 hours, tissues were cryoprotected in 30% sucrose for three days, then embedded in O.C.T Compound (Electron Microscopy Sciences) and sectioned at 50 µm thickness using a CM3050 cryostat (Leica Microsystems). Imaging was performed with a STELLARIS 8 confocal microscope (Leica Microsystems) equipped with a HC PL APO 63x/1,40 OIL CS2 oil immersion objective (Leica Microsystems). A montage of multiple Z-stacks of 180×180×30μm size with 2μm z-intervals was taken to image the whole tissue in high resolution. Excitation and emission were done in three channels: in the DiD range (Ex: 646 nm, Em: 656-750nm), for detecting the liposomes, and in two background channels, which were above and below the DiD range (Ex: 485 nm, Em: 490-600 nm; Ex: 705 nm, Em: 710-840 nm). ImageJ was used for quantification, with background subtracted from DiD range and the remaining DiD signal/ image field quantified.

### In vivo biodistribution, retention, and endothelial interaction of liposomal formulations

Retention and clearance of liposomal formulations were assessed in 5-month-old male C57BL6 mice. Groups included untreated controls (N=3), FL/RSV-treated (N=3), and ApoE-FL/RSV-treated (N=3) mice. The treatment groups received daily retro-orbital injections of 30 µL of 10 mg/mL DiD-labeled liposome solutions for four consecutive days, while control mice received no injection. Mice were sacrificed 30 minutes after the final injection on the fourth day. To determine whether liposomes interact with the cerebrovascular endothelium directly VECAD Rosa26-tdTomato mice (9 months old, tamoxifen-induced at 3 months) were divided into three groups: untreated control (N=3), FL/RSV (N=3), and ApoE-FL/RSV (N=3). The treatment groups received retro-orbital injections of 120 µL of 10 mg/mL DiR-labeled liposome solution for two consecutive days, while control mice received no injection. Mice were sacrificed on the fourth day, and tissues were processed. Detailed images of organs were obtained across three channels: tdTomato (Ex: 555 nm, Em: 565-700 nm), DiR (Ex: 748 nm, Em: 760-850 nm), and background (Ex: 640 nm, Em: 650-700 nm). Z-stack of 485×485×50 μm size with 1 μm z-intervals was acquired. For whole-organ imaging HC PL APO 10x/0,40 CS2, objective (Leica Microsystems) was used to generate Z-stacks of 50 μm with 5 μm z-intervals from the DiR channel (Ex: 748 nm, Em: 760-850 nm). Maximum Intensity Z projection was used for visualization and ImageJ was used to subtract the background and adjust brightness and contrast in both the tdTomato and DiR channels. Native images in orthogonal view were used for qualitative evaluation.

### Functional ultrasound imaging for the assessment of deep cerebral vasculature and neurovascular coupling

#### Plastic cranial window surgery

Cranial window surgeries were performed as described above (“Cranial window surgeries”), except that a custom-cut curved plastic window^33^ (TPX™, thickness 0.125 mm, Mitsui Chemicals) was used in place of the glass window. The window was secured to the skull using Starbond GAP FILLER super glue (Amazon), followed by the application of a polymerization accelerant. All other surgical steps, including anesthesia, scalp incision, craniotomy, and postoperative care, were identical to those described before^132^.

#### Functional ultrasound imaging

Two weeks post-surgery, the animal was lightly anesthetized (1-1.5% isoflurane), intubated, and positioned on a thermoregulated stereotactic frame (51625W, Stoelting Co, Wood Dale, IL, USA), while attached to a SomnoSuite® Low-Flow Anesthesia System (Kent Scientific Corporation, Torrington, CT, USA) to control for changes in respiratory patterns that could affect hemodynamic stability. The mouse’s head was shaved with a hair removal cream (Nair body cream with Aloe) and cleaned with EtOH. The ultrasonic probe (IcoPrime-4D MultiArray 15 MHz, ICONEUS, France) of the ICONEUS One functional ultrasound device (ICONEUS, France) was positioned directly above the cranial window and submerged in ultrasound gel (Gel de contact, Drexco Medical, France). Functional ultrasound data was captured using the scanner’s live acquisition software (IcoScan, Iconeus, Paris, France). All fUS data were acquired using a set of 10 tilted plane-wave transmissions (Transmit frequency: 15 MHz, Pulse widths: 2 cycles per pulse, -12° to +12° tilting angles) compounded with a 5000 Hz pulse repetition frequency (PRF) leading to a 500 Hz ultrasound imaging frame rate. Raw data for each of the four linear arrays were beamformed independently in the receive mode and independently summed coherently in the transmit mode. Each subarray (4×64 elements) is tightly spaced with a 2.1 mm spacing between subarrays, to optimize the field of view while minimizing crosstalk. The four images were simultaneously obtained from the corresponding four linear arrays at 500 Hz. Each imaging session lasted 1.5–2 hours. Subsequently, the animal was closely observed until it fully emerged from anesthesia, ensuring the absence of any abnormal behavior or signs of discomfort. Ultrasound Localization data was captured using the scanner’s live acquisition software (IcoScan, Iconeus, Paris, France). Transmit voltage was limited to 8V (mechanical index (MI) of 0.1) to prevent microbubble destruction during ULM. All ULM data was acquired using a set of 5 tilted plane-wave transmissions (Transmit frequency: 15 MHz, Pulse widths: 2 cycles per pulse, -12° to +12° tilting angles) compounded with a 5000 Hz pulse repetition frequency (PRF) leading to a 1000 Hz ultrasound imaging frame rate for a better estimation of higher speeds in larger arteries. ULM Raw data for each of the four linear arrays were beamformed independently in the receive mode and independently summed coherently in the transmit mode.

#### Microbubble suspension injection

ULM, also known as Super-Resolution Ultrasound Localization Microscopy, involves detecting and tracking individual microbubbles (MBs) injected intravenously into the animal. In this study, we injected 50 µL of sterile MB suspension (DEFINITY, Lantheus, Billerica, MA, USA) to capture high-definition images of microbubble trajectories within a 500 µm thick coronal brain volume. Before the retro-orbital injection, DEFINITY MBs were activated by vigorous agitation for 45 seconds until the clear solution turned milky. This technique, combined with the transparent TPX window and a two-week recovery period post-surgery, allowed for high-resolution imaging (5µm/pixel) from the brain’s surface to its base.

#### Ultrafast ultrasound imaging parameters and image generation

Image acquisition was carried out using the ICONEUS One ultrasound system (Iconeus One – 256 channels, ICONEUS, France), equipped with a 15MHz MultiArray transducer (IcoPrime-4D MultiArray, ICONEUS, France). Successive raw ultrasound images were recorded for 10 minutes at a rate of 1 kHz, each image representing the coherent sum of 8 different transmission angles (-12°, -8.57°, -5.14°, -1.71°, 1.71°, 5.14°, 8.57°, 12°) at an 8 kHz pulse repetition frequency. A bolus of MBs was injected at the start of each 10-minute recording session.

Using the "Compute SuperLoc" software integrated into the ICONEUS One fUS imager (ICONEUS, France), we generated a maximum projection of all tracked microbubble trajectories within a single coronal brain section. This projection covered the entire 10-minute recording following MB injection. The "Display SuperLoc" software was then used to extract key data, including an MB intensity map, a microbubble speed map, and an MB direction map. These raw data outputs formed the basis of our analysis, used to derive cerebrovascular physiological endpoints.

#### ULM image analysis

All image processing and analysis were conducted using FIJI ImageJ v. 1.52p software, except for RGB velocity image conversion from hexadecimal color code to grayscale by interpreting each 6-digit hexadecimal color code as a 24-bit base-16 number and converting it into its decimal equivalent, producing a single scalar intensity value for each pixel, which was performed using MATLAB R2023a. Exported high-resolution ULM recordings were initially processed. Images from each measurement were stacked, and batch segmentation was performed using the Weka Trainable Segmentator, a supervised machine learning algorithm in ImageJ, to generate binary vascular masks. The final step involved assessing total vascular area density using binary images from the cortical and hippocampal regions as 2D representations of the 3D VOI.

## Statistical analysis

If not otherwise stated, all data are presented as means ± standard error of the mean (SEM). GraphPad Prism 9 Software (Dotmatics, La Jolla, CA, USA) was employed for the statistical analyses. Comparisons between groups of experimental results were conducted using T-test, repeated T-test, one-way ANOVA with Fisher LSD post hoc test or an equivalent non-parametric (Kruskal-Wallis) or non-equal distribution (Brown-Forsythe), or Repeated-Measure ANOVA tests whichever was applicable based on the sample type and distribution. A minimum of three independent measurements were performed for all data (n≥3), and the exact "N" animal numbers are specified in the figure legends where it is applicable. Levels of significance were denoted as follows: * p<0.05, ** p<0.01, *** p<0.001. Data points affected by an animal’s condition which was not part of the experimental protocol or e.g. low quality of its cranial window were excluded from the study, to reduce disturbing external factors.

## Reporting summary

Further information on research design is available in the Nature Portfolio Reporting Summary linked to this article.

## Data availability statement

The data supporting the findings of this study can be found either in the main text or can be obtained from the authors upon a reasonable request. For access to this data, please reach out to the corresponding authors. The mass spectrometry proteomics data have been deposited to the ProteomeXchange Consortium via the PRIDE partner repository [PubMed ID: 39494541] with the dataset identifier PXD072125 (reviewer access token: ATUG9BBhLxId).

## Code availability

All software used for acquisition or data analysis is either openly or commercially available. The custom-made ImageJ macros for blood-brain barrier permeability measurement, *in vitro* colocalization experiment, are in Supplementary File 1,2.

## Acknowledgements

We extend our heartfelt appreciation to the team at the Division of Comparative Medicine, University of Oklahoma Health Sciences Center, for their exceptional support in animal care and for sharing their profound expertise with us. A special note of thanks goes to Dr. Shawn Lane, DVM, whose invaluable guidance and proficiency have been crucial in both surgical and postsurgical care phases. We also thank Dr. Wendy Williams, DVM, for her pivotal role in designing effective pre- and post-surgical treatment protocols. Our gratitude further extends to Ms. Carlye Yancey, BS, for her outstanding expertise in animal husbandry. Our acknowledgement would be incomplete without expressing our deep gratitude to Ms. Julie Farley for her meticulous execution of spatial learning and memory experiments within the Animal Behavioral Core of OUHSC, and to Ms. Tripti Gautam for her diligent efforts in preparing scRNA-seq samples. We are equally thankful to the Data Science and Me biology data science company for their exemplary services in single-cell sequencing analysis and data visualization, contributing significantly to the success of this research. S.B. gratefully acknowledges Philipp Schönnenbeck for his help with the data analysis in CryoVIA. The authors gratefully acknowledge the electron microscopy training, imaging and access time granted by the life science EM facility of the Ernst-Ruska Centre at Forschungszentrum Jülich as well as the computing time granted by the JARA Vergabegremium and provided on the JARA Partition part of the supercomputer JURECA at Forschungszentrum Jülich (http://dx.doi.org/10.17815/jlsrf-7-182). The 4.0 version of ChatGPT, developed by OpenAI, was used as a language tool to refine our writing, enhancing the clarity of our work.

This work was supported by grants from the American Heart Association (RG: AHA916225, ANT: 24CDA1276505, and ST: AHA CDA941290), the Oklahoma Center for the Advancement of Science and Technology, the National Institute on Aging (RF1AG072295, R01AG055395, R01AG068295; R01AG070915, K01AG073614, K01AG073613, R03AG070479, R21AG080775-01A1), the National Institute of Neurological Disorders and Stroke (R01NS100782), the National Cancer Institute (R01CA255840), the Oklahoma Shared Clinical and Translational Resources (U54GM104938) with an Institutional Development Award (IDeA) from NIGMS, the Presbyterian Health Foundation, the Reynolds Foundation, the Oklahoma Nathan Shock Center (P30AG050911), the NCI Cancer Center Support Grant (P30 CA225520) and the Oklahoma Tobacco Settlement Endowment Trust and Hevolution Grant (HF-GRO-23-1199084-16). The project was also supported by the Ministry of Innovation and Technology of Hungary from the National Research, Development and Innovation Fund, financed under the TKP2021-NKTA funding scheme; by funding through the National Cardiovascular Laboratory Program (RRF-2.3.1-21-2022-00003) provided by the Ministry of Innovation and Technology of Hungary from the National Research, Development and Innovation Fund; Project no. 135784 implemented with the support provided from the National Research, Development and Innovation Fund of Hungary, financed under the K20 funding scheme and the European University for Well-Being (EUniWell) program (grant agreement number: 101004093/ EUniWell/EAC-A02-2019 / EAC-A02-2019-1). BC was supported by the EKÖP-2025-703 founding scheme through University Research Scholarship Programme of the Ministry of Culture and Innovation from the source of the National Research, Development and Innovation Fund. The funding sources played no part in the design of the study, the collection, analysis, and interpretation of data, the writing of the report, or the decision to submit the article for publication. The authors bear sole responsibility for the content, which may not necessarily reflect the official views of the National Institutes of Health, the American Heart Association, or the Presbyterian Health Foundation.

## Author contributions

The foundational concept of this study was developed by AnC and AgC, while the overall design and interpretation of data were collaboratively contributed to by all authors. AgC and TG have established the liposome preparation protocol for the experiments, did the liposome quality check and the further characterization. PM contributed as a protein corona expert to characterize the physicochemical parameters of the liposomes.TG, PH and MK carried out MS studies, SB, CS cryo-EM analysis. SS, TG, NH, RP, PS, SC, RG, KVK have carried out the *in vitro* and related experiments, including cell culturing, treatment, sorting, and microscopy analysis. BC and SS have established liposomal preparation protocol for *in vivo* studies. BC, ANT, DN, LH, MS, RYN, RK, EGB, PM, AY, ZB have assessed the *in vivo* treatments, surgeries and experiments and analyses. Neurovascular coupling assessments were carried out by SSC, SN, and ST. HA, WJ, TGO, TK contributed to sequencing data analysis and interpretation. The initial manuscript draft was co-authored by ANT, AnC, AgC, RM and ZU, with all authors actively participating in the manuscript’s revisions. This collective effort culminated in an unanimously approved final manuscript.

## Conflict of interest statement

The underlying technology is patent pending in Europe (PCT/EP2025/075652). The patent is owned by the RE3B Therapeutics UG. AgC and NH hold shares in this company. The remaining authors declare no competing interests.

## Disclosure of financial interests

The authors declare no competing financial interests.

## Abbreviations

AAV: Adeno-associated virus
ACN: Acetonitrile
AMPK: AMP-activated protein kinase
ANOVA: Analysis of variances
ApoE: Apolipoprotein E
a.u.: Arbitrary unit
AUC: Area under curve
BBB: Blood-brain barrier
BCA: Bicinchoninic acid
CBF: Cerebral blood flow
CNS: Central nervous system
Cryo-TEM: Cryogenic transmission electron microscopy
CRISPR: Clustered regularly interspaced short palindromic repeats
DiO: 3,3’-dioctadecyloxacarbocyanine perchlorate
DiD: 1,1’-dioctadecyl-3,3,3 ’,3 ’-tetramethylindodicarbocyanine 4-chlorobenzenesulfonate
DiR: 1,1’-dioctadecyl-3,3,3 ’,3 ’-tetramethylindotricarbocyanine iodide
DLS: Dynamic light scattering
DOPE: 1,2-dioleoyl-sn-glycero-3-phosphoethanolamine
DOTAP: 1,2-dioleoyl-3-trimethylammonium-propane
EL: Endocytic liposomes
ELS: Electrophoretic light scattering
ER: Endoplasmic reticulum
EtOH: Ethanol
eNOS: endothelial nitric oxide synthase
FA: Formic acid
FITC: Fluorescein isothiocyanate
FL: Fusogenic liposomes
GFAP: Glial fibrillary acidic protein
GFP: Green fluorescent protein
GO: Gene ontology
hCMEC/D3: Human cerebral microvascular endothelial cells, D3 cell-line
HEPES: 4-(2-hydroxyethyl)-1-piperazineethane-sulfonic acid
HP: Human plasma
IBA1: Ionized calcium-binding adaptor molecule 1
iBAQ: Intensity-based absolute quantification
IL6: Interleukin-6
KD: Knockdown
LC-MS/MS: Liquid chromatography-tandem mass spectrometry
LDLR: Low-density lipoprotein receptors
LRP1: Low-density lipoprotein receptor-related protein 1
MP: Mouse plasma
NeuN: Neuronal nuclei
NF-κB: Nuclear factor kappa B
NO: Nitric oxide
NVC: Neurovascular coupling
PBS: Phosphate buffered saline
PFA: Paraformaldehyde
PO-RSV: Per os-resveratrol
RAWM: Radial arm water maze
riBAQ: Relative intensity-based absolute quantification
RIPA: Radioimmunoprecipitation assay buffer
RSV: Resveratrol
RT: Room temperature
RT-qPCR: Quantitative reverse transcription polymerase chain reaction
scRNA-seq: Single-cell RNA sequencing
SIRT1: Sirtuin 1
SP3: Solid phase 3
TNF-α: Tumor necrosis factor α
ULM: Ultrasound localization microscopy
VCID: Vascular cognitive impairment and dementia
VECAD: Vascular endothelial cadherin
VOI: Volume of interest
WB: Western blotting
WGA: Wheat Germ Agglutin

---

**Extended Data Figure 1.**
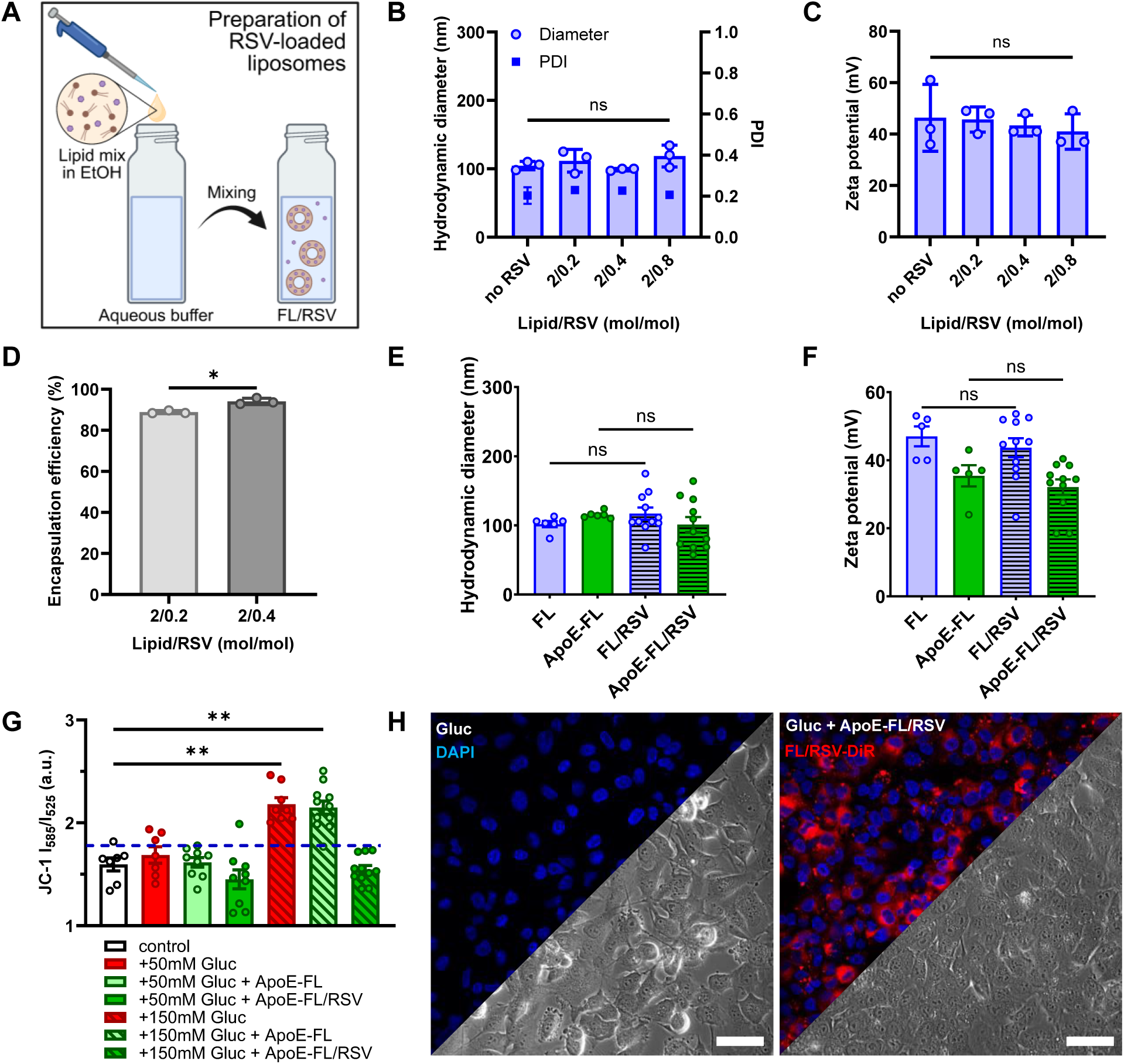
Physical characteristics of FL/RSV with and without ApoE corona as well as pharmacological effect of ApoE-FL/RSV treatment on mitochondrial condition *in vitro*. A) FL can be easily prepared by pipetting the ethanol solution of the lipid mixture DOPE/DOTAP/aromatic molecules (2/2/0.2 mol/mol) in an aqueous buffer. B) Hydrodynamic diameter, polydispersity index (PDI) and C) zeta potential of FL containing RSV (FL/RSV) at different concentrations were determined using dynamic and electrophoretic light scattering, respectively. Particle characteristics did not change significantly upon RSV loading, indicating complete RSV incorporation into the liposomes. D) RSV encapsulation efficiency (EE) into FL was determined via particle centrifugation and subsequent quantification of unbound RSV using fluorescence spectroscopy. EE values exceeded greater than 80% in all cases. E–F) Characterization of FL and FL/RSV with and without APOE corona showed no impact of RSV loading on particle size and zeta potential. G) ApoE-FL/RSV treatment led to mitochondrial membrane potential recovery after glucose-induced hyperpolarization, thus maintaining mitochondrial function and preventing elevated ROS production as well as apoptotic pathways. H) Representative micrographs of hyperglycemic hCMEC/D3 cells with and without ApoE-FL/RSV treatment (Scale bars, 50 µm). Data are mean±SEM. Statistical significance was calculated using two-sample t-test and is indicated by *p<0.05, **p<0.01, ***p<0.001.

**Extended Data Figure 2.**
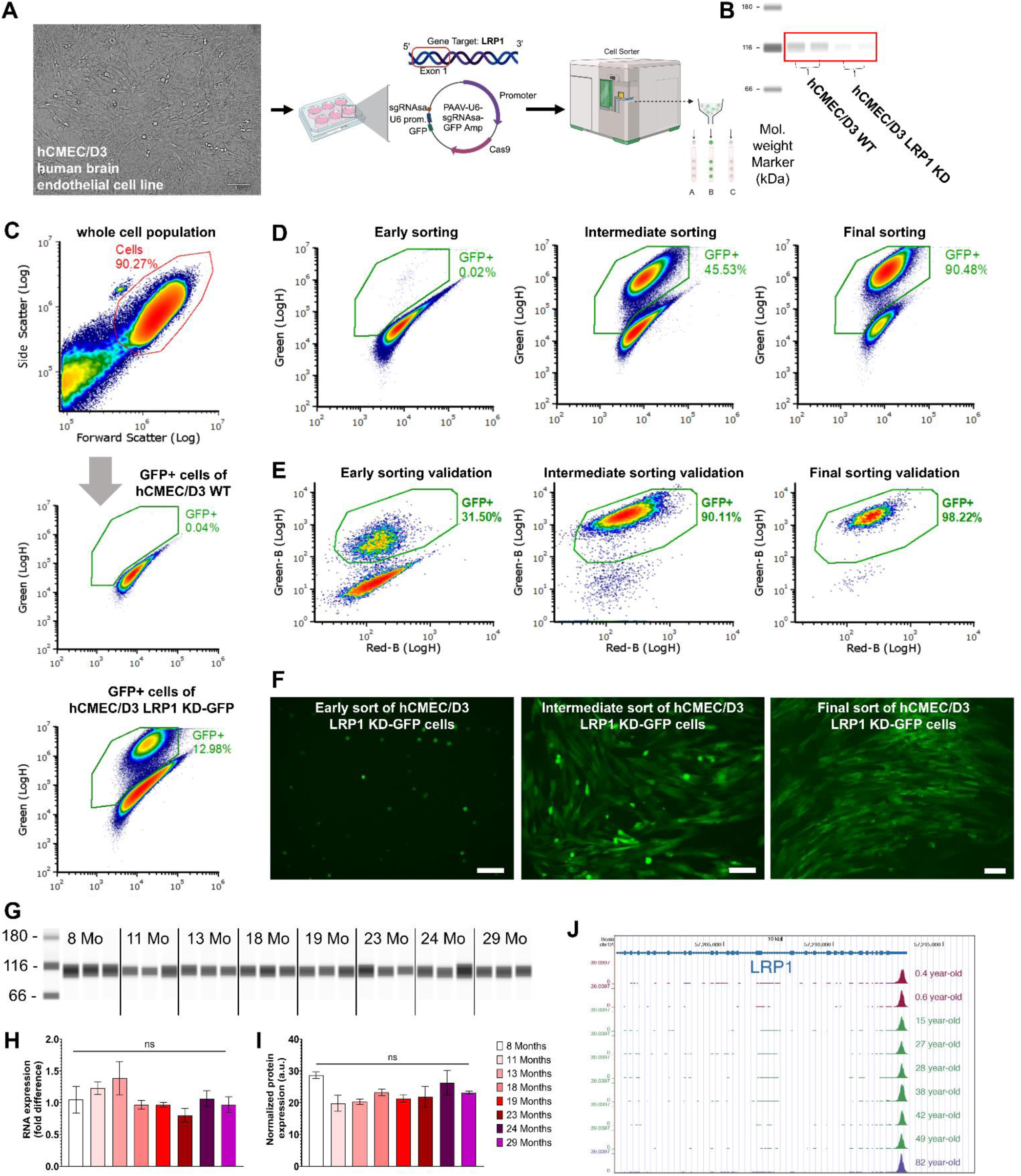
Establishment of LRP1-KD hCMEC/D3 for examining the connection between LRP1 and ApoE in liposome docking. **A)** Brightfield image of the human brain endothelial cells before transfection (Left). These cells were then treated with PAAV-U6-sgRNAsa-GFP Amp vector in 6-well plates and incubated for 72 hours (Middle). After 72 hours, LRP1-KD, GFP-positive cells were repeatedly sorted until the maximum knockdown efficiency was achieved. **B)** Confirmation of LRP1 knockdown in sorted cells. LRP1 KD was verified using an automated western blotting instrument (JESS). **C)** Gating strategy for sorting the LRP1-KD, GFP-positive hCMED/D3 cells. The population of hCMEC/D3 cells was selected based on forward and side scatter (Top). Wild-type hCMED/D3 cells (middle) were used to determine the GFP-positivity gate. GFP-positive cells (Bottom) were sorted to establish a pure population of LRP1-KD hCMEC/D3 cells. **D**–**E)** Representative flow cytometry dot plots from early, intermediate, and final stages of the sorting process. These dot plots demonstrate progressive enrichment in LRP1-KD-GFP-positve cells, during **(D)** and after **(E)** each round of sorting. **F)** Representative images of LRP1-KD GFP-positive cells from all stages of sorting. **G)** Expression pattern of LRP1 in the brain tissue of mice of different ages using Western blot. **H)** Quantification of LRP1 mRNA expression shows no changes during aging. **I)** Similarly, LRP1 protein expression shows no significant changes in aging mice. **J)** Furthermore, our analysis of publicly available scRNA-seq datasets shows that LRP1 expression is maintained in aging human cerebrovascular endothelial cells as well. Data are mean±SEM, n≥3 for all groups. *p<0.05, **p<0.01, ***p<0.001 with ANOVA test, and Tukey post-hoc.

**Extended Data Figure 3.**
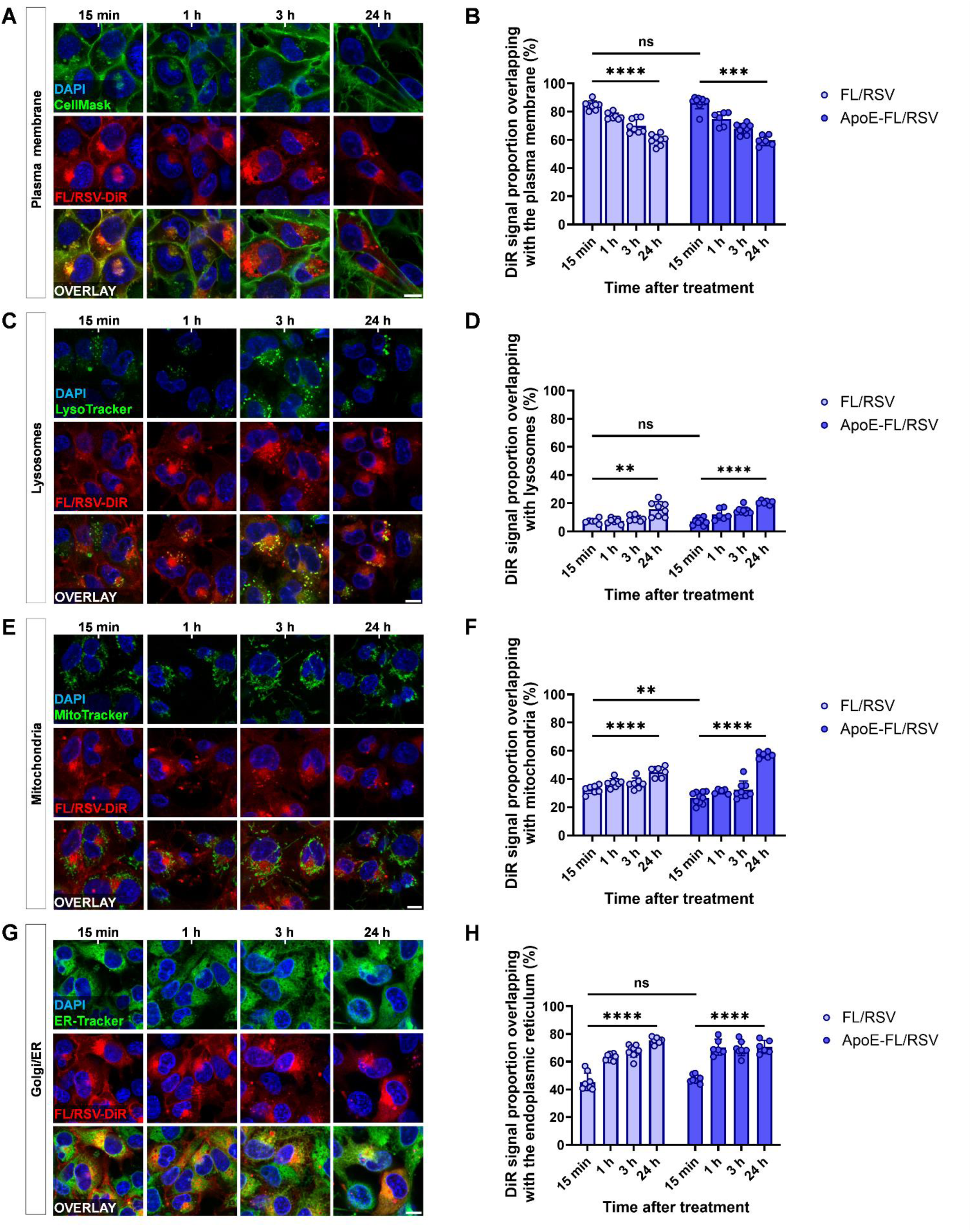
Time-dependent intracellular localization of FL/RSV-DiR with and without ApoE targeting motive in different cellular compartments. Representative fluorescence microscopy images of hCMECs at various time points after FL/RSV treatment accompanied by bar charts representing the DiR proportion overlapping with the fluorescence staining of the plasma membrane (A–B), lysosomes (C–D), the mitochondria (E–F), and the ER/golgi apparatus (G–H). Nuclei were labeled with DAPI (blue), whereas cell organelles were stained in green by applying CellMask, LysoTracker, MitoTracker, and ER-Tracker, respectively. The extent of colocalization was determined by measuring the pixel ratio of the overlapping DiR signal to the total DiR signal. All scale bars correspond to 10 µm.

**Extended figure 4.**
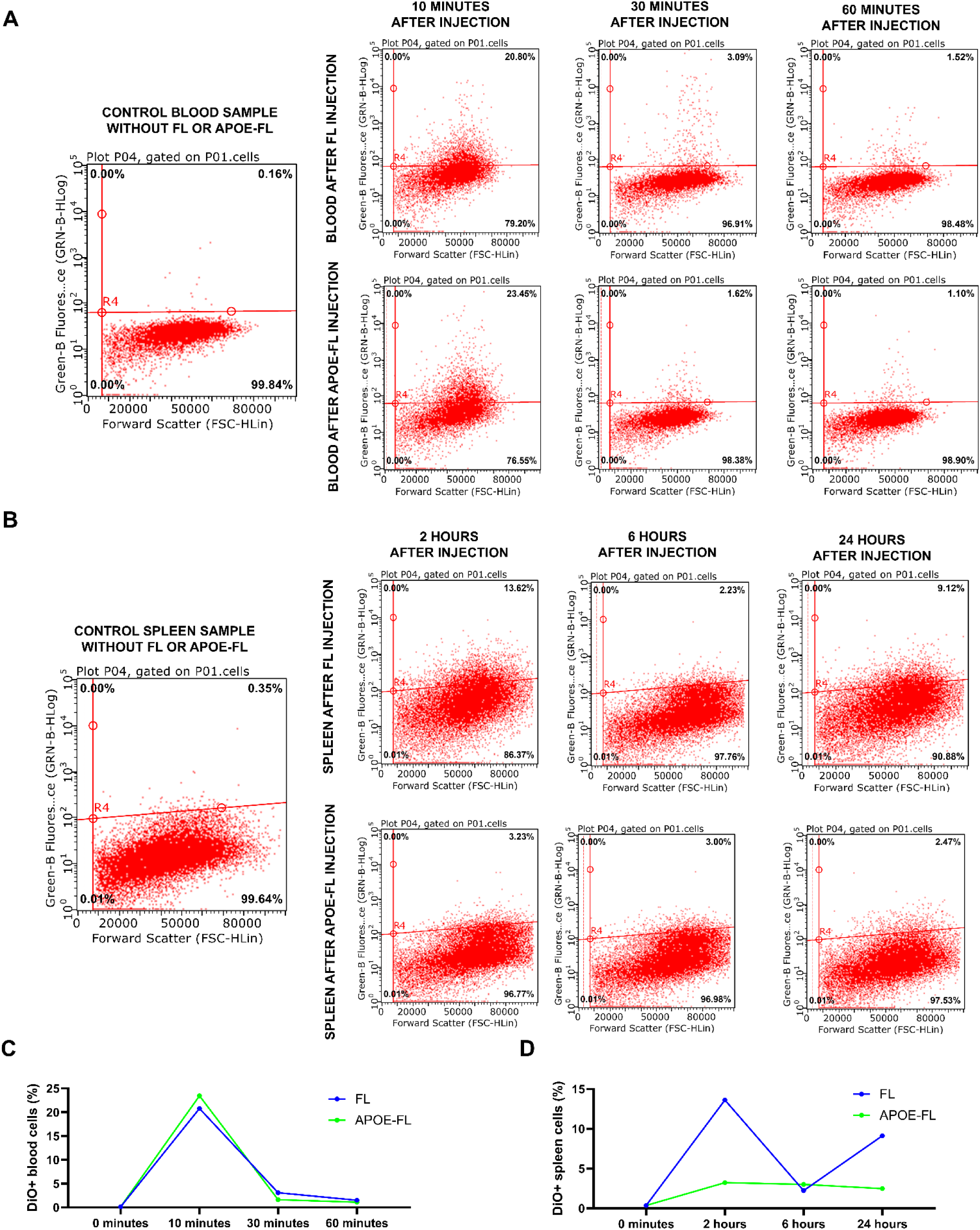
Transient interaction of fusogenic liposomes with circulating immune cells and minimal uptake by splenic immune populations. A) Flow cytometry analysis of DiO-associated fluorescence in circulating blood cells following retro-orbital injection of FL-DiO or ApoE-FL-DiO. A control blood sample without liposomes shows negligible background fluorescence. After FL injection, a transient increase in DiO⁺ events is detected at 10 minutes, consistent with short-lived liposome attachment to circulating white blood cells. This signal rapidly diminishes by 30 minutes and is nearly absent by 60 minutes. ApoE-FL-DiO shows a similar transient pattern but with slightly fewer DiO⁺ circulating cells overall. No evidence of sustained intracellular uptake was detected in either formulation. B) Flow cytometry analysis of splenic immune cells at 2, 6, and 24 hours after injection of FL-DiO or ApoE-FL-DiO. Control spleen samples show minimal autofluorescence. A modest increase in DiO⁺ splenic cells is observed at 2 hours exclusively in the FL-treated group, whereas ApoE-FL shows no appreciable accumulation at any time point. By 6 and 24 hours, DiO⁺ splenic events approach baseline for both formulations. C) Quantification of DiO⁺ circulating blood cells over time. Both FL and ApoE-FL exhibit a sharp, transient rise in DiO-associated events at 10 minutes, followed by rapid clearance by 30 minutes. D) Quantification of DiO⁺ splenic cells over time. A small but detectable increase is present only in the FL group at 2 hours, with negligible uptake in the ApoE-FL group. Both groups return to baseline by 24 hours.

**Extended Data Figure 5.**
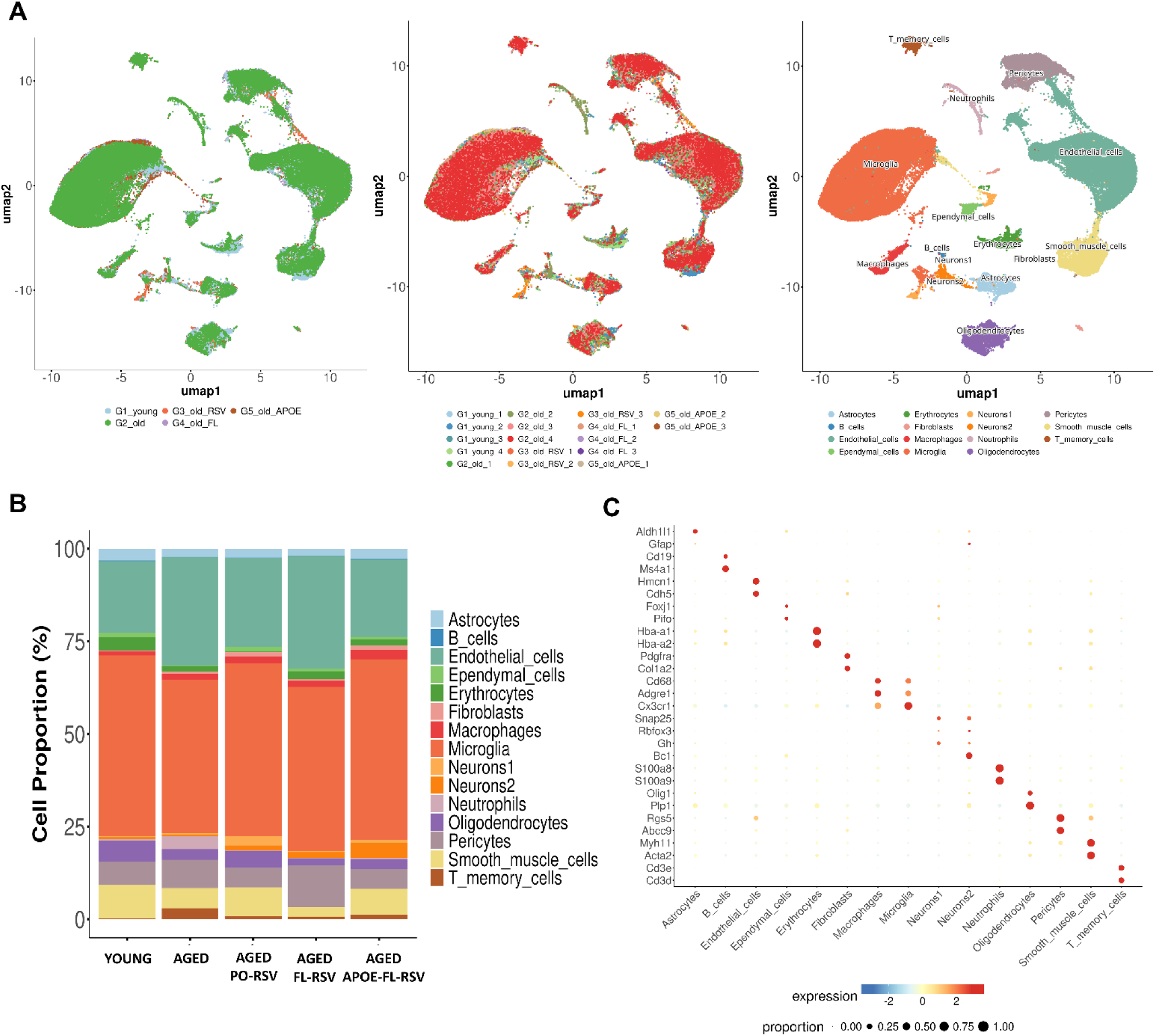
Single-cell RNA-seq analysis of brain cell populations following aging and nanoparticle treatments. **A)** UMAP visualization of all single-cell transcriptomes from young, aged, and aged mice treated with *per os* (PO)-RSV, FL-RSV, or ApoE-FL-RSV. **Left:** Samples grouped by experimental condition. **Middle:** Overlay of treatment-specific cell distributions showing the relative contribution of each experimental group. **Right:** Unsupervised clustering annotated by major brain cell types, including neurons, astrocytes, oligodendrocytes, microglia, endothelial cells, pericytes, smooth muscle cells, macrophages, ependymal cells, fibroblasts, erythrocytes, neutrophils, and T-memory cells. **B)** Stacked bar plots showing the proportional abundance of major brain cell types across conditions. **C)** Dot plot displaying canonical marker genes used to assign cell-type identities. Dot color indicates scaled expression level, and dot size represents the proportion of cells expressing each marker within a given cluster. These markers confirm the robustness of cell-type annotations across neuronal, glial, vascular, and immune lineages.

**Extended Data Figure 6.**
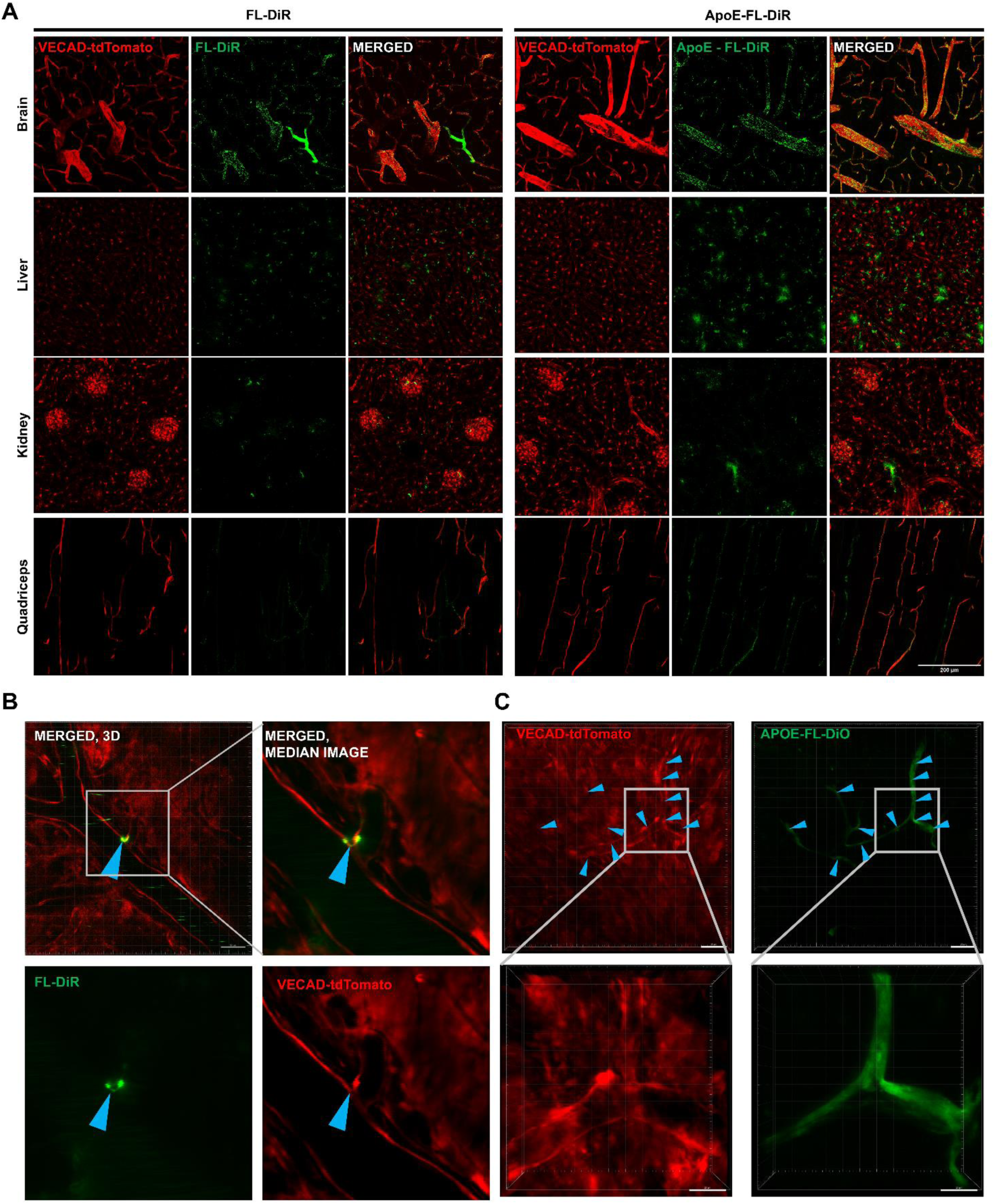
Organ-Level biodistribution and cerebrovascular endothelial colocalization of FLs in VECAD-tdTomato mice. **A)** Representative images about the *in vivo* accumulation of fluorescent signal originating from FL-DiR and ApoE-FL-DiR in the brain, liver, kidney, and skeletal muscle tissues of the VECAD-tdTomato mice. The scale bar represents 200μm. **B)** FL-DiR visualized after docking to tdTomato expressing brain endothelium. Cerebrovasculature has been recorded as a TPM as a t-stack and in different time points and visualized with IMARIS software. Scale bar represents 40μm. **C)** ApoE-FL-DiO visualized after docking to tdTomato expressing brain endothelium. Cerebrovasculature has been recorded as a TPM as a t-stack and in different time points and visualized with IMARIS software. The scale bar represents 40μm.

**Extended Data Figure 7.**
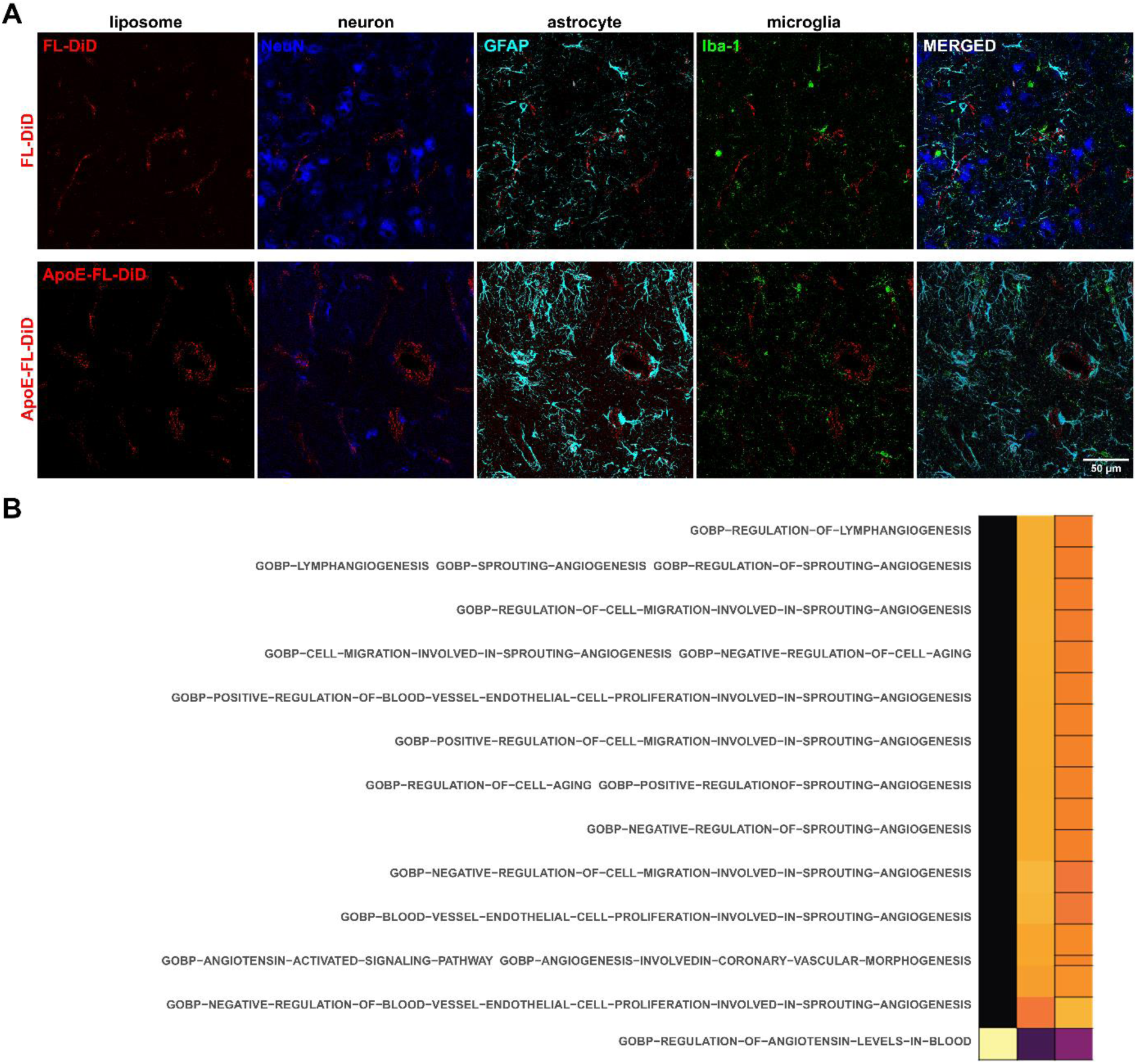
Cellular localization and gene ontology analysis of FL-DiD and ApoE-FL-DiD Nanoparticles in Mouse Brain Tissue. **A)** Representative confocal microscopy images demonstrating cellular localization of FL-DiD and ApoE-FL-DiD particles in mouse brain tissue. Images illustrate distribution of lipid dye DiD in neurons (NeuN, blue), astrocytes (GFAP, cyan), and microglia (Iba-1, green), with merged images indicating spatial overlap and interaction of nanoparticles with various cell types. Scale bar: 50 µm. **B)** Heatmap showing Gene Ontology Biological Process (GOBP) terms significantly enriched in response to nanoparticle treatment. Enriched pathways include angiogenesis, endothelial cell migration, proliferation, and processes related to vascular regulation and cellular aging. Each row represents an individual GOBP term, with color intensity indicating the significance of enrichment.

**Extended Data Figure 8.**
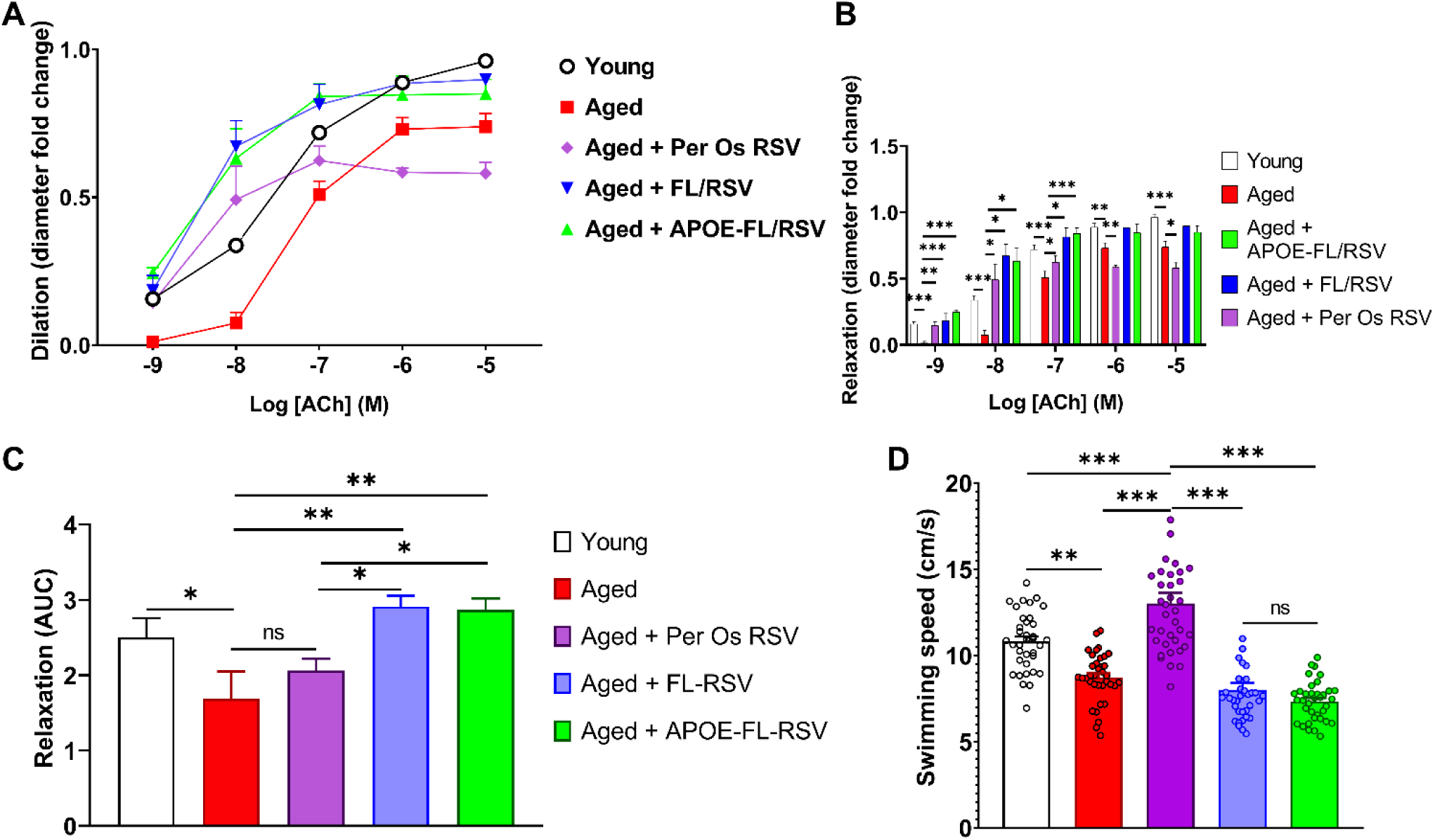
FL-RSV and ApoE-FL-RSV improve aged animal aorta function. **A)** Dilation curves of the aorta rings during acetylcholine-induced vasorelaxation in mice. A significant difference in dilation between groups was observed when 4μM acetylcholine (ACh) was applied to aorta rings. **B)** Quantification of aorta ring muscle relaxation in response to Ach. Data are mean±SEM. *p<0.05, **p<0.01, ***p<0.001 (n≥3 for all groups) with Kruskal-Wallis test. **C)** Summarized quantification of aorta rings relaxation in response to Ach with AUC (area under curve). Data are mean±SEM. *p<0.05, **p<0.01, ***p<0.001 (n≥3 for all groups) with ANOVA, Fisher LSD post-hoc test. **D)** Average swimming speeds in the RAWM for each group, analyzed with One-way ANOVA, revealed no significant differences across in response to injected FL-RSV or ApoE-FL-RSV, underscoring that swimming speed did not influence learning performance. On the other hand, the *per os* RSV showed a significant increase in swimming speed without an improvement of cognitive performance. Data are presented as mean±SEM (n=10–15 per group). Data are mean±SEM (n=10–15 per group). Statistical significance is indicated by *p<0.05, **p<0.01, ***p<0.001 using ANOVA.

